# Transcriptome-wide mapping of small-molecule RNA-binding sites in live cells

**DOI:** 10.1101/2024.05.30.596700

**Authors:** Yuquan Tong, Patrick R. A. Zanon, Xueyi Yang, Xiaoxuan Su, Jessica L. Childs-Disney, Matthew D. Disney

## Abstract

Small molecules targeting RNA can be valuable chemical probes and potential therapeutics. The interactions between small molecules, particularly fragments, and RNA, however, can be difficult to detect due to their modest affinities and short residence times. Here, we describe the procedures for mapping the molecular fingerprints of small molecules in vitro and throughout the human transcriptome in live cells, identifying both the targets bound by the small molecule and the sites of binding therein. For complete details on the use and execution of this protocol, please refer to ^1^.

## Introduction

The essential roles of RNA are well-documented in every aspect of life.^2^ Small molecules that selectively bind to RNA targets can be leveraged as chemical tools to probe RNA structure and function, as well as lead molecules to rescue disease-associated cellular phenotypes.^3, 4^ The discovery of these small molecules, however, can be challenging, as the interactions between small molecules and RNA targets can be difficult to detect, particularly in cells. One method that addresses this challenge is Chemical Cross-Linking and Isolation by Pull-down (Chem-CLIP),^5^ where a covalent bond is formed between the small molecule and the RNA target, capturing a dynamic binding event. This method has been successfully used for target validation and for mapping of small molecule binding sites.^6, 7^ Since its first report in literature,^5^ other methods have been coined but share the same underlying principles of Chem-CLIP to study target engagement via the formation of covalent bonds between RNA and small molecules, including PEARL-seq^8^ and reactivity-based RNA profiling (RBRP).^9, 10^

Here, we describe the protocol for our recent platform advances, including an optimized *in vitro* method, a method to create molecular fingerprints across the human transcriptome in live cells, and a method to map small molecule binding sites transcriptome-wide. Chem-CLIP probes (Figure 1) comprise an RNA-binding small molecule appended with a cross-linking module (a diazirine) and a pull-down handle (an alkyne). The probes are incubated with RNA targets either *in vitro* or in live cells and form covalent bonds with bound targets upon UV irradiation; that is, reversible binding interactions are captured by irreversible photoaffinity labeling. The modified RNAs are then immobilized and enriched by copper-catalyzed azide-alkyne cycloaddition (CuAAC)^11, 12^ to disulfide-linked azide beads (Figures 2 and 3). For transcriptome-wide studies, RNAs are eluted from the beads and subjected to transcriptome-wide RNA-seq analysis. Since the cross-linked small molecule can impede reverse transcription,^7^ the termini of the truncated complementary DNA (cDNA) can be used to map the binding sites of the small molecule to its bound RNA targets.

**Figure 1.**
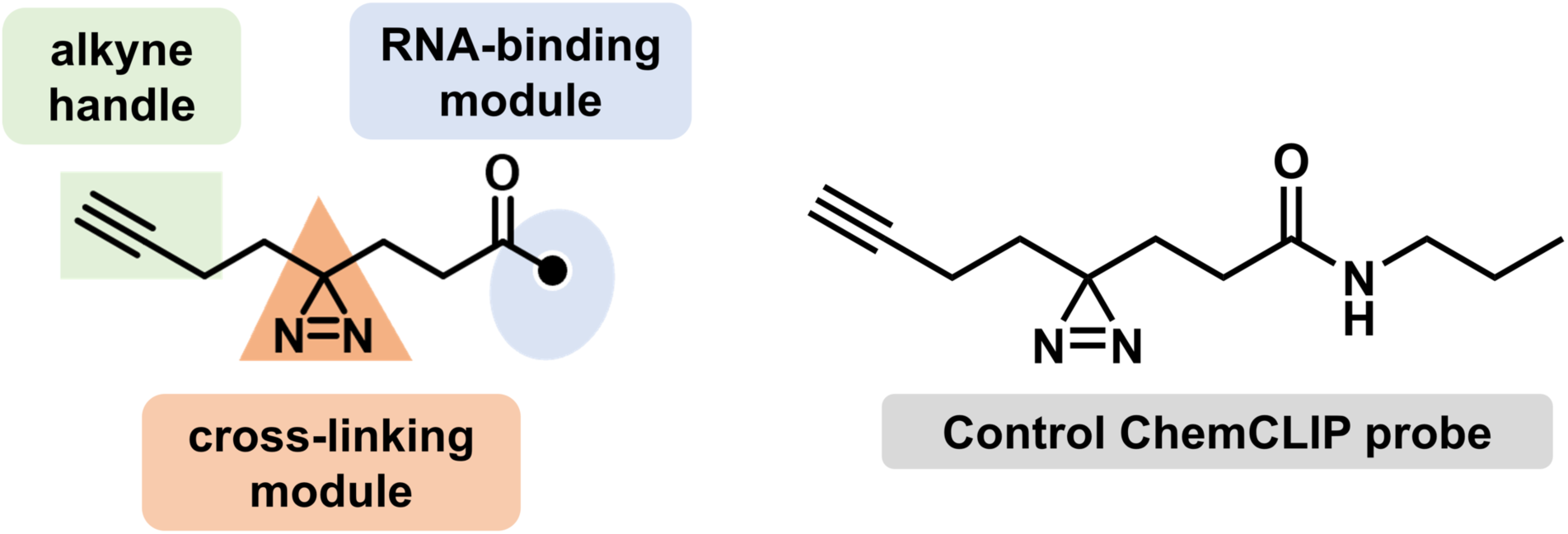
The general structures of a Chem-CLIP probe and a control probe that lack a potential RNA-binding module. The Chem-CLIP probe consists of an RNA- binding module tethered to a diazirine cross-linking module and an alkyne handle. As the RNA-binding module drives target engagement, the diazirine cross-linking module can be photoactivated under UV light to react with RNA targets and form covalent bonds. The biorthogonal alkyne enables click reaction with azide-functionalized beads or azido-biotin. Placement of the diazirine and alkyne handles should be informed by structure-activity relationship studies to ensure that functionalization does not affect molecular recognition.

**Figure 2.**
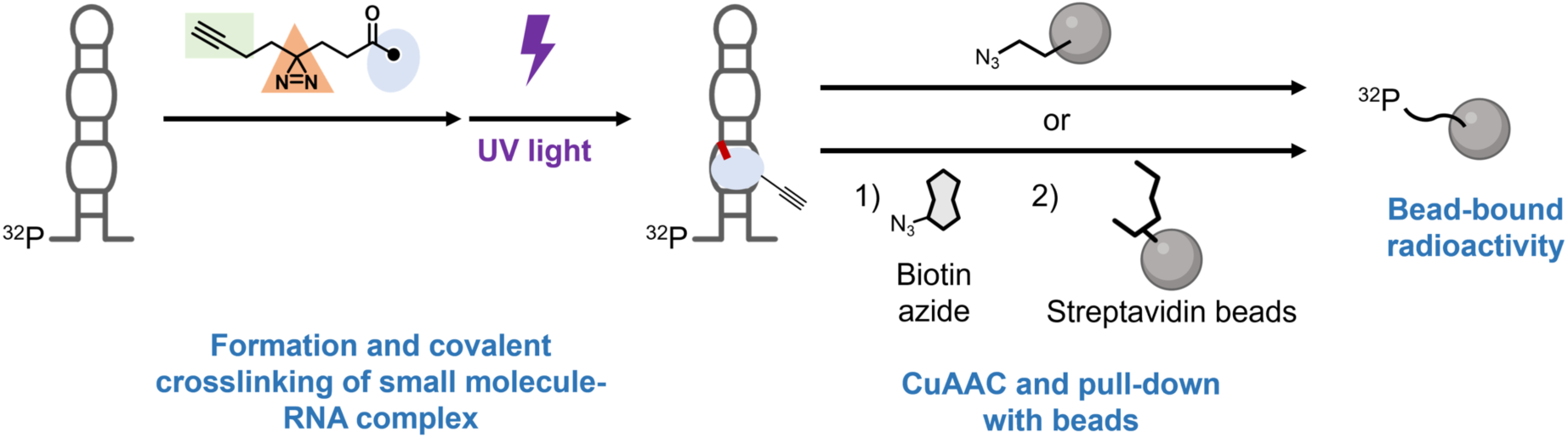
Target occupancy studies by *in vitro* Chem-CLIP (Part I). To study target occupancy/engagement *in vitro*, a radiolabeled (or fluorescently labeled) RNA of interest is incubated with a Chem-CLIP probe. Following UV irradiation, the covalent complex is “clicked” either directly onto azide beads or first to biotin azide, which is then captured by streptavidin beads. After stringent washing, the now bead-bound radioactivity is quantified by scintillation counting to measure the fraction of RNA that interacts with the Chem-CLIP probe. Note, Chem-CLIP studies, whether conducted *in vitro* or in cells, can also be performed as a competition experiment to study a parent RNA-binding small molecule.

**Figure 3.**
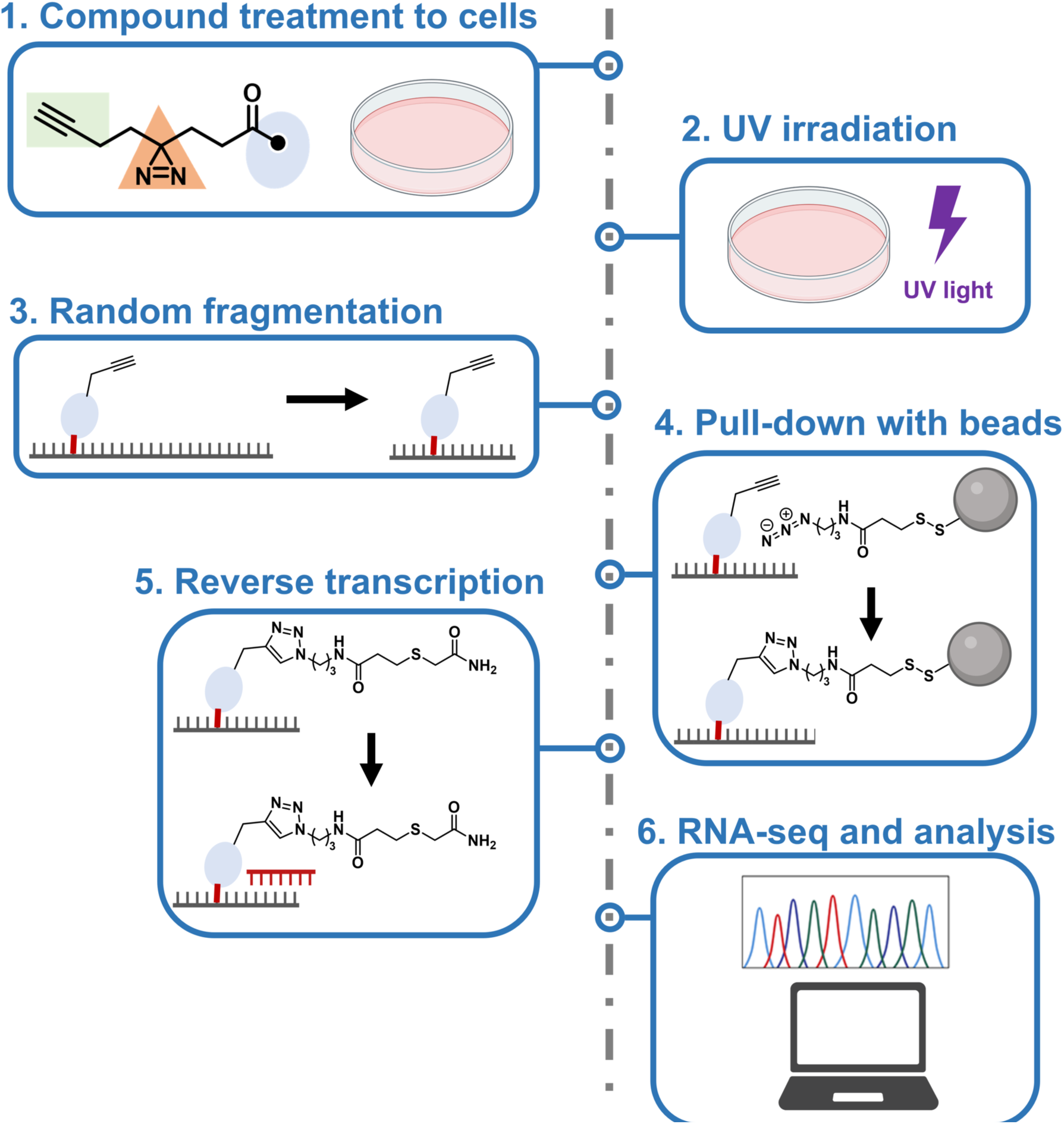
Schematic of the Chem-CLIP workflow for transcriptome-wide mapping of small molecule-RNA binding sites in cells (Part II). In brief, cells are treated with a Chem-CLIP probe, typically at 20 mM for 16 h. To induce reaction of the Chem-CLIP probe with bound RNAs, that is formation of a covalent bond, the cells are irradiated with UV light and total RNA is harvested. The RNA is then fragmented and the fragments that have been cross-linked to the Chem-CLIP probe are isolated/purified with azide-functionalized beads by a click reaction with the biorthogonal alkyne handle. Bound targets and the binding sites within them are identified by RNA-seq analysis as described in the main text.

The protocol below describes the specific steps for: (i) target occupancy studies *in vitro*; (ii) creating transcriptome-wide ligandability maps in MDA-MB-231 human triple- negative breast cancer (TNBC) cells; and (iii) mapping the binding sites of small molecules in cells. The latter two protocols can also be applied to other cell lines.

### [Preparation of buffers and reaction solutions] Timing: [2 h]

1. Prepare the following buffers in Nanopure water and store at room temperature. The pH of buffers is adjusted by using either 1 M NaOH or 1 M HCl.

a. 1× HEPES Buffer (25 mM HEPES, pH 7.0, where HEPES is 4-(2- hydroxyethyl)-1-piperazine ethanesulfonic acid)
b. 1× Wash Buffer (10 mM Tris-HCl, pH 7.0, 4 M NaCl, 1 mM ethylenediaminetetraacetic acid (EDTA), 0.1% (v/v) Tween-20)
c. 3 M sodium acetate, pH 5.2
2. Prepare the following stock solutions separately in Nanopure water. All solutions must be prepared fresh prior to each experiment.

a. 10 mM CuSO4·5H2O, pH ∼4
b. 250 mM sodium ascorbate, pH ∼7
c. 50 mM THPTA (tris(3-hydroxypropyltriazolylmethyl)amine), pH ∼7-8
d. 200 mM TCEP (tris(2-carboxyethyl)phosphine hydrochloride), pH ∼3
e. 100 mM K2CO3, pH ∼11
f. 200 mM iodoacetamide

**CRITICAL**: All buffers and solutions must be prepared by using RNase-free water.

Key resources table

| REAGENT or RESOURCE | SOURCE | IDENTIFIER |
| --- | --- | --- |
| Cell line and culturing reagents |  |  |
| MDA-MB-231 cells | ATCC | HTB-26 |
| RPMI 1640 | Corning | 10-041-CV |
| Fetal Bovine Serum (FBS) | Sigma Aldrich | F2442 |
| Penicillin/Streptomycin Solution, 50× | Corning | 30-001-CI |
| 0.25% Trypsin-EDTA, phenol red | Gibco | 25200-056 |
| 1× Dulbecco's Phosphate-buffered Saline (DPBS), without calcium and magnesium | Corning | 21-031-CV |
| **Chemicals** |  |  |
| THPTA (tris(3-hydroxypropyltriazolylmethyl)amine) | Vector Laboratories | CCT-1010 |
| TCEP (tris(2-carboxyethyl)phosphine) hydrochloride | Sigma Aldrich | 51805-45-9 |
| Iodoacetamide | Sigma Aldrich | 144-48-9 |
| Copper(II) sulfate pentahydrate | Sigma Aldrich | 7758-99-8 |
| Potassium carbonate | Sigma Aldrich | 584-08-7 |
| [γ-<sup>32</sup>P]-ATP | Revvity | BLU035C001MC |
| Acrylamide | Acros Organics | 164850025 |
| Bis-acrylamide | RPI International | A11270 |
| Urea | Oakwood Chemical | 044699 |
| Tris base (2-Amino-2-(hydroxymethyl)-1,3-propanediol) | RPI International | T60040 |
| Boric acid | Sigma-Aldrich | B0252 |
| Ethylenediaminetetraacetic acid (EDTA) disodium salt | EMD Millipore | 324503 |
| Ammonium persulfate (APS) | Sigma-Aldrich | A3678 |
| Tetramethylethylenediamine (TEMED) | Sigma-Aldrich | T9281 |
| THPTA (tris(3-hydroxypropyltriazolylmethyl)amine) | Click Chemistry Tools | 760952-88-3 |
| Copper(II) sulfate pentahydrate | Sigma-Aldrich | 7758-99-8 |
| Biotin-PEG3-azide | Sigma-Aldrich | 762024 |
| 1× Dulbecco's Phosphate-buffered Saline (DPBS), without calcium and magnesium | Corning | 21-031-CV |
| Sodium ascorbate | RPI International | S42175 |
| Sodium dodecyl sulfate (SDS) | Fisher Scientific | BP166 |
| Sodium deoxycholate | Thermo Scientific | J62288.22 |
| Nonidet P40 substitute | Thermo Scientific | J19628-AP |
| **Critical commercial kits or assays** |  |  |
| Quick-RNA Miniprep Kit | Zymo Research | R1054 |
| Disulfide Azide Agarose beads | Vector Laboratories | CCT-1238 |
| NEBNext Magnesium RNA fragmentation module | New England Biolabs | E6150S |
| RNAClean XP beads | Beckman Coulter | A66514 |
| NEBNext rRNA Depletion Module | New England Biolabs | E6310 |
| NEBNext Ultra II Directional RNA Library Prep Kit for Illumina | New England Biolabs | E7760 |
| T4 Polynucleotide Kinase | New England Biolabs | M0201L |
| Azide Magnetic Beads | Vector Laboratories | CCT-1036 |
| Dynabeads MyOne Streptavidin C1 | Invitrogen | 65002 |
| **Deposited data** |  |  |
| RNA-seq data by using this method | Mendeley Data | doi:10.17632/56r9zmjps2.1 |
| **Software and algorithms** |  |  |
| STAR | <sup>13</sup> | version 2.7.9 |
| Genrich | github.com/jsh58/Genrich | version 0.6.1 |
| bamCompare | <sup>14</sup> | version 3.5.0 |

### Part I. Step-by-step method details for *in vitro* Chem-CLIP with radiolabeled RNA

Radiolabeling of the RNA

### Timing: [1 h]

1. In a 0.6 mL microcentrifuge tube, add the following:
2. 10 pmol RNA (e.g. 10 µL of 1 µM stock)
3. 1.5 µL of 10× T4 PNK buffer
4. 1 µL of T4 polynucleotide kinase (10,000 U/mL)
5. DNase/RNase-free water so that the total volume in Step 1 is 14.6 µL
6. Behind the plexiglass shield, preferably in a fume hood, carefully add 0.4 µL of [γ-

### 32P]-ATP (6000 Ci/mmol, 150 mCi/mL)

***Note***: All experiments that use radioactive materials must follow the safety guidelines of the respective institution. Alternatively, a fluorescently labeled RNA could be used, although this experimental protocol has not been applied to such RNAs.

a. 3. Incubate sample at 37 °C for 45 min Purification of radiolabeled RNA **Timing: [5 h]**
b. 4. Gel electrophoresis
c. Prepare a denaturing polyacrylamide gel. The following will afford 50 mL of 15% (w/v) polyacrylamide gel suitable for a gel with following dimensions: 18 cm wide, 15 cm high, and 2 mm thick, holding a 20-well comb (where each well is 1.7 cm deep and 5 mm wide).
d. 30 mL of 25% acrylamide (23.75% w/v acrylamide, 1.25% w/v bis-acrylamide, 7.5 M urea)
e. 5 mL of 10× Sequagel Buffer (890 mM Tris, pH 8.4, 980 mM boric acid,

40 mM EDTA)

a. iii) 15 mL of diluent (7.5 M urea)
b. iv) 500 µL of 10% (w/v) ammonium persulfate in water (APS), 40 µL of tetramethylethylenediamine (TEMED)
c. Dilute samples with 2× Loading Dye (2 mM Tris, pH 8.0, 8 M urea, 20 mM EDTA, and 0.02% (w/v) orange G) and separate the radiolabeled RNA from unincorporated [γ-^32^P]-ATP by electrophoresis for 1:40 h at 25 W, using 1× TBE (130 mM Tris, pH 7.6, 45 mM boric acid, 2.5 mM EDTA) as the running buffer.
d. 5. Scan and extract RNA from gel.
e. Carefully separate the plates and cover the gel in its entirety with plastic wrap.
f. Expose a phosphor screen to the gel, typically 5 min or less, at −20 °C.
g. Image by phosphorimaging, for example using a Typhoon FLA9500 (GE Healthcare), print, and use the resulting template to excise the radiolabeled RNA. The gel can be imaged again to confirm successful excision of the radiolabeled RNA.
h. Place the excised gel containing the radiolabeled RNA in a 2 mL centrifuge tube and add 400 µL of 300 mM NaCl. Place the tube in a tube rotator for 3-5 h at 4 °C.
i. Transfer supernatant to a fresh 1.5 mL microcentrifuge tube, being careful not to transfer the gel pieces.
j. Add 1 mL of 20 mg/mL glycogen (optional) followed by 1 mL EtOH and place the sample at −80 °C for 2 h.
k. Centrifuge the sample for 15 min at 12,000 × g and 4 °C.
l. Carefully remove the EtOH without disturbing pellet.
m. Place the tube on its side and let dry inside a fume hood for up to 30 min or place in a vacuum concentrator for 2 min to remove residual EtOH.
n. Resuspend residue in 50 µL of Nanopure water.
o. 6. Transfer 1 µL of the purified RNA into a 0.6 mL microcentrifuge tube and quantify the RNA’s concentration by liquid scintillation counting, for example using a Beckman Coulter LS 6500 Multi-Purpose Scintillation Counter.

Pull-down of radiolabeled RNA by photoaffinity probes

### Timing: [estimated 6 h for 12 samples]

a. 7. In a 1.5 mL microcentrifuge tube (Axygen Maxymum Recovery; cat# MCT-150-L-C), fold RNA (≥6000 cpm per replicate) in appropriate Reaction Buffer (e.g. 1× DPBS).
b. e.g. for r[GCG(CUG)12CGC], prepare the RNA in 1× DPBS, place at 95 °C for 1 min and then immediately place on ice for ≥ 10 min.

***Note***: To control for unintended or non-specific interactions, compounds that engage the target should be tested against appropriate control RNAs that lack the desired binding site. In the present example, a construct with A-U base-pairs instead of 1×1 nucleotide UU loops would fulfill this purpose.

a. b) The volume required per condition for technical triplicates is 65 µL. Therefore, to study 10 photoaffinity probes at a single concentration, fold 715 µL of RNA (this includes 10% excess to account for errors in pipetting). Please note additional volume will be required for controls listed in (c) and that, while, subsequent replication of results is recommended, independent replicates can also be included directly in this experiment.

***Note***: As for any RNA-small molecule interaction, the choice of buffer can have a significant impact on the observed pull-down. Experiments are typically carried out in triplicates, however, duplicates may be better suited for screening, for example a library of fully functionalized fragments (FFFs).

a. c) To ensure the validity of the experiment, include the following control samples:
b. No treatment/enrichment – serves as a reference for the input
c. DMSO – serves as reference for non-specific pull-down of the RNA
d. No UV – without UV irradiation, no covalent cross-link should be established, ablating pull-down
e. No “click” – without the copper catalyst, no enrichment should be observed
f. 8. Aliquot 65 mL of the folded RNA into siliconized 0.6 mL microcentrifuge tubes (G-Tube, BIO PLAS, cat# 4160SL), keeping an aliquot (65 mL to afford three technical triplicates of 20 mL each) that will not be further processed as reference in technical triplicates to measure the total amount of radioactivity added to each sample (No treatment/enrichment listed in (7c)).
g. 9. Add 0.65 µL 100× compound stock in DMSO to RNA, affording a final DMSO concentration of 1% (v/v).

***Note***: Concentrations of 100 µM or less are recommended to avoid potential aggregation and precipitation of the photoaffinity probe and/or the probe-RNA complex.

a. 10. Immediately after addition of the photoaffinity reagent, transfer technical replicates (20 µL) to fresh, siliconized 0.6 mL microcentrifuge tubes.
b. 11. Incubate the samples for 30 min in the dark.
c. 12. Irradiate the samples with 350-365 nm UV light for 15 min in a photo-crosslinker (UV Stratalinker 2400 with Eiko F15T8/BL bulbs). Ensure that the centrifuge tube lids are open.
d. 13. While waiting for the irradiation to be completed, place 10 µL of Azide Magnetic Beads (10 mg/mL, 30-50 nmol azide/mg) per sample (3-5 nmol azide) into a 1.6 mL microcentrifuge tube. Wash the beads with 2× volumes of Reaction Buffer three times, centrifuging (1,000× g for 10 s) and removing the supernatant between each wash.

***Note:*** The amount of beads can be reduced depending on the experiment, however, a minimum 1.5-fold excess over the alkyne probe is recommended. Alternatively, this step can be performed by using Streptavidin-functionalized beads. For this alternative procedure, please see *Pull-down with Streptavidin beads* at the end of this section.

a. 14. Resuspend the beads in Reaction Buffer (2.2× original volume of beads used in step 13)
b. 15. Prepare a click reaction master mix consisting of following amounts per sample. Note 10% excess should be prepared to account for errors in pipetting.
c. 1 µL of 10 mM CuSO4 in water
d. 0.6 µL of 50 mM THPTA in water (freshly prepared)
e. 0.6 µL of 250 mM sodium ascorbate in water (freshly prepared). The pH of the sodium ascorbate should be ∼7-8.
f. 20 µL of washed Azide Magnetic beads from Step 14
g. 16. Add 22.2 µL of this mix to all samples, making sure to pipet up and down prior to dispensing to ensure equal distribution of the beads.
h. 17. Place the tubes on their sides on a rotating platform (250 rpm) and incubate samples for 1 h at 37 °C, ensuring that the beads stay in suspension and do not settle.
i. 18. Briefly spin down samples (1,000× g for 10 s).
j. 19. Add 300 µL of 1× DPBS containing 0.1% (w/v) SDS. Mix well by pipetting up and down.
k. 20. Wash the beads.
l. Place samples in a magnetic stand.
m. Once the solution is cleared of the beads, transfer the supernatant to a 5 mL scintillation vial where all washes of a particular sample will be collected.
n. Add 300 µL of 0.1% (w/v SDS) in 1× DPBS to the beads and mix by pipetting.

***Note:*** If more samples are being processed than the number that can be accommodated by the magnetic rack, add wash buffer to the beads before processing additional samples.

a. d) Shake samples with tubes on the side for 10 min as described in Step 17.
b. e) Briefly spin down samples (1,000× g for 10 s).
c. 21. Repeat step 20 twice with High Salt Wash Buffer (1× DPBS, 850 mM NaCl, 0.1% (w/v) SDS, 0.5% (w/v) sodium deoxycholate, 1% (v/v) Nonidet P40 substitute
d. 22. Repeat step 20 twice with Low Salt Wash Buffer (1× DPBS, 150 mM NaCl, 0.1% (w/v) SDS, 0.5% (w/v) sodium deoxycholate, 1% (v/v) Nonidet P40 substitute.
e. 23. Place samples in magnetic stand.
f. 24. Once cleared, carefully transfer the supernatant into its respective scintillation vial.
g. 25. Add 200 µL of Low Salt Wash Buffer, pipet up and down, and transfer beads to a new scintillation vial, keeping the pipet tip affixed to the pipet to use as described in Step 26.
h. 26. Use a different pipet to add 200 µL of Low Salt Wash Buffer to the now-empty microcentrifuge tube. With the pipet + tip used in Step 25, transfer to the scintillation vial containing the beads to ensure full transfer.
i. 27. Put the now emptied and rinsed microcentrifuge tube into new scintillation vial.
j. 28. Measure the radioactivity present in all fractions (beads, supernatant, empty tubes) for each sample by scintillation counting.

### Alternative route at Step 13: Pull-down with Streptavidin beads

a. 13) Prepare a click reaction master mix consisting of following amounts per sample.

Note 10% excess should be prepared to account for errors in pipetting.

a. 1 µL of 10 mM CuSO4 in water
b. 0.6 µL of 50 mM THPTA in water (freshly prepared)
c. 0.6 µL of 250 mM sodium ascorbate in water (freshly prepared). The pH of the sodium ascorbate should be ∼7-8.
d. 1.2 µL of 1 mM biotin-PEG3-azide (Sigma-Aldrich, CAS #875770-34-6, cat# 762024); i.e., 1.2 nmol (vs. alkyne probe 50 µM / 1 nmol)
e. 14) Add 3.4 µL of the click reaction master mix to each sample (except the No “click” control described in Step 7).
f. 15) Incubate the samples for 1 h at 37 °C.
g. 16) Add 40 µL of Dynabeads MyOne Streptavidin C1 (as a slurry, directly from the bottle without washing), pipetting up and down immediately. The capacity of 40 µL of Dynabeads MyOne Streptavidin C1 is 400 pmol oligonucleotide or >1.1 nmol free biotin.

***Note***: For probe concentrations higher than 50 µM, the amounts of biotin-PEG3-azide and streptavidin beads must be adjusted accordingly. Ethanol precipitation after CuAAC can remove excess unbound probe and biotin.

1. 17) Place the tubes on their sides on a rotating platform and incubate the samples at 37 °C for 30 min with shaking at 250 rpm.
2. 18) Follow steps 20-28 as described above.

Data analysis

For the analysis of the enrichment, the average radioactivity detected in “no treatment/enrichment” samples serve as reference of complete enrichment, *i.e.* 100% capture of radioactivity on the beads. Accordingly, the relative enrichment for a given sample is determined as follows:

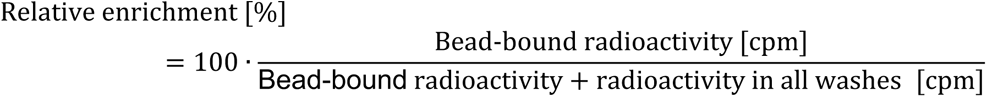

The total sum of radioactivity across fractions (beads, supernatant/washes, tubes) should be consistent between enriched samples and not be less than ∼90% of the “no treatment/enrichment” references.

The DMSO-treated sample controls for non-specific interactions of the RNA with the beads, tubes, and pipet tips. The enrichment observed should not exceed 5% and, while some low-level photo-activation might occur due to ambient light, the “no UV” controls should be approximately the same as this vehicle background.

### Part II. Step-by-step method details for transcriptome-wide mapping in live cells

Maintaining and splitting cells

### Timing: [1-3 days]

MDA-MB-231 cells are maintained in an incubator at 37 °C with 5% CO2. Cells from cryo-stocks must be passaged at least two times before performing experiments, and cells should be discarded when the passage number exceeds 20.

1. Recover MDA-MB-231 cells from cryogenic storage.
2. Prepare growth medium by supplementing RPMI 1640 medium with 10% (v/v) fetal bovine serum (FBS) and 1× Penicillin/Streptomycin.
3. Warm ∼13 mL of the medium to 37 °C. Add 12 mL to a 100 mm diameter dish.
4. Warm the cryo-stock at 37 °C, add 1 mL of pre-warmed medium, and then transfer to the 100 mm diameter dish containing the pre-warmed growth medium.
5. Gently rotate the dish and then place in an incubator overnight.
6. Replace medium with pre-warmed, fresh growth medium the second day.

***Note:*** Different types of cells may need different handling procedures and growth medium. Please refer to the vendor’s information for other cell types/lines.

i. 2. Maintain and sub-culture MDA-MB-231 cells
ii. Change the growth medium every 2-3 days by aspirating the old growth medium and then adding 12 mL of pre-warmed, fresh growth medium. Subculture the cells when they reach >90% confluency, as monitored by microscope.
iii. To subculture the cells, remove the growth medium with a serological pipet and rinse the cells with 2 mL of 1× DPBS.
iv. Add 2 mL of 0.25% Trypsin-EDTA and place the dish in the incubator for

∼3 min. Monitor detachment of the cells by microscope. Once they begin to detach, proceed immediately to (d).

i. d. Add 4 mL of fresh growth medium and transfer all of the medium containing the detached cells to a 15 mL sterile conical tube.
ii. e. Centrifuge at 160 × g for 2 min and carefully remove the supernatant with a serological pipet. Resuspend the cells in 8 mL growth medium. Transfer 2 mL of the resuspended cells in a new 100 mm diameter dish containing 10 mL of pre-warmed growth medium (1:4 ratio) and transfer to an incubator. When subcultured in this fashion, the confluency of the 100 mm diameter dish will be ∼40% after overnight incubation.

**CRITICAL**: Cells must be checked for mycoplasma contamination before experiments commence. We use PromoKine PCR Mycoplasma Test Kit (PK-CA91-1024) for this purpose.

Compound treatment & RNA extraction

### Timing: 1 day

In the following steps, live cells are treated with a Chem-CLIP probe of fully functionalized fragment and cross-linked to bound targets. T cellular RNA is then harvested and isolated, including treatment with DNase I.

i. 3. Cells are appropriate for compound treatment upon reaching ∼80-90% confluency.
ii. From 20 mM compound stock solutions, prepare a 1:1000 dilution with fresh growth medium (10 mL per 100 mm diameter dish) such that the final concentration is 20 µM (0.1% v/v DMSO).
iii. Remove the old growth medium with a serological pipet and add the freshly prepared growth medium containing the compound. For each compound, two 100 mm diameter dishes are treated in parallel as biological duplicates.
iv. The following control groups should be performed in parallel (each with a minimum of two biological replicates), including (1) a control diazirine probe lacking the RNA-binding module; (2) a vehicle treated group (no compound, 0.1% (v/v) DMSO; and (3) a compound-treated group without UV-cross-linking.
v. Incubate the cells in an incubator for 16 h.
vi. 4. After 16 h, remove the growth medium using a serological pipet and rinse cells with 2 mL of 1× DPBS.
vii. 5. Add 2 mL of 1× DPBS to the cells and expose cells under UV light (UV Stratalinker 2400) for 10 min at room temperature. Ensure that the lid is removed during irradiation.

***Note***: Cells are cross-linked in 1× DPBS to maintain viability.

**CRITICAL:** Ensure that the lid is removed during irradiation.

i. 6. Carefully remove the 1× DPBS by pipet and extract total RNA by using a Quick-RNA Miniprep Kit (Zymo; R1054) per the manufacturer’s recommended protocol as follows:
ii. Add 1.4 mL of Lysis Buffer directly to the 100 mm diameter dish and incubate at room temperature for 2 min.

***Note***: Gently swirl the dish to make sure the Lysis Buffer covers the entire surface of the dish. Other RNA extraction kits/methods may be used depending on the types of samples.

**CRITICAL**: All tubes and pipette tips must be free of RNase.

i. b. Transfer the cell lysate to two yellow Spin-Away Filters columns (700 µL each) placed into collection tubes and centrifuge at 12,000 × g for 1 min. Keep the flow-through as it contains the total RNA. Discard the columns.
ii. c. Add 700 µL of 100% ethanol directly to the flow-through in the collection tubes. Mix by pipetting.
iii. d. Place green Zymo-Spin IIICG columns onto new collection tubes. Transfer 700 µL of the mixture from the previous step to the columns and centrifuge at 12,000 × g for 1 min. Discard the flow-through and repeat for the remaining mixture.
iv. e. Add 350 µL of Wash Buffer to the column and centrifuge at 12,000 × g for 1 min. Discard the flow-through.
v. f. Prepare the DNase I digestion solution by adding 5 µL of DNase I per column to 75 µL of Digestion Buffer per column and mix by pipetting up and down (include 10% excess). Add 80 µL of this solution to each column and incubate the samples at room temperature for 15 min.

***Note***: DNase I treatment is necessary to remove genomic DNA, which will interfere with the downstream RNA-seq analysis.

a. g. Add 400 µL of Prep Buffer to the column and centrifuge at 12,000 × g for 1 min. Discard the flow-through.
b. h. Add 400 µL of Wash Buffer to the column and centrifuge at 12,000 × g for 1 min. Discard the flow-through.
c. Add 700 µL of Wash Buffer to the column and centrifuge at 12,000 × g for 2 min. Transfer the column to a clean 1.6 mL microcentrifuge tube.

***Note***: When transferring the column, ensure that the bottom of the column does not come into contact with the flow-through to avoid contamination.

a. j. Add 50 µL of RNase-free water to the column and incubate at room temperature for 2 min. Centrifuge at 12,000 × g for 1 min.
b. k. Combine the eluted RNA from the two columns. Note in Step 6b that the lysate was split into two columns per sample.
c. 7. Quantify the eluted RNA and assess its quality by measuring A260/A280 with a NanoDrop spectrophotometer, for example a Thermo Fisher NanoDrop 2000. Samples with a A260/A280 ratio of 2.00 ± 0.15 are considered to be of high enough quality to proceed. Values deviating from this indicate contamination,

e.g. from residual gDNA or insufficient washing.

***Note***: For one 100 mm diameter dish of MDA-MB-231 cells, total RNA concentration is expected to be around 80-150 ng/µL when eluted in a total of 100 µL water (combining the 50 µL elutions from the two columns used per sample).

**Pause point**: RNA samples can be stored at −20 °C for 1 week or at −80 °C for a month. For samples stored longer than these periods, the integrity of RNA should be assessed by using a Bioanalyzer or gel electrophoresis prior to proceeding to the next steps.

Random fragmentation of RNA

### Timing: 6 h

The cross-linked RNA (now containing the alkyne group from the compound) is randomly fragmented to ∼150 nucleotide (nt) lengths. This allows for the pull-down of only regions cross-linked by the compound instead of the entire transcript. It should be noted that fragmentation can be performed either before or after the pull-down, each with distinct advantages and disadvantages. Fragmentation *before* pull-down provides specific mapping of the binding sites but is not suitable for transcripts with low abundance or a large number of repeats. Complementarily, fragmentation *after* pull-down allows for high read coverage as it enriches the entire transcript; however, small molecule binding sites cannot be mapped.

a. 8. Total RNA is randomly fragmented to ∼150 nt lengths by using an NEBNext Magnesium RNA fragmentation module (New England Biolabs, E6150S). The minimum amount of total RNA used as input in this step is 5 µg, although 10 µg or more is recommended. The following steps are performed per manufacturer’s recommendations.
b. For every 100 µL of harvested RNA, add 10 µL RNA Fragmentation Buffer (10×) and mix by pipetting.
c. Incubate at 95 °C for 3 min.
d. Immediately place the tube on ice and allow it to cool down for 1 min. Centrifuge for 30 s by using a benchtop mini centrifuge (2000 × g).
e. Add 10 µL of RNA Fragmentation Stop solution (10×) and mix by pipetting.
f. 9. If 100 µL of RNA was used in Step 8a, then add 12 µL of 3 M sodium acetate, pH 5.2, and 400 µL of 100% ethanol. If larger volumes of RNA were used, scale the amount of sodium acetate and ethanol accordingly. Mix by pipetting and place in −80 °C for at least 4 h.

**Pause point**: The solution can also be left at −80 °C overnight.

a. 10. Centrifuge at >12,000 × g at 4 °C for 20 min. Carefully remove the supernatant by pipetting.

***Note***: The precipitated RNA pellet may not be visible to the naked eye. Gently tilt the tube and avoid disturbing the pellet by carefully pipetting out liquid from the side opposite from where the pellet has formed.

a. 11. Gently add 400 µL of 70% (v/v) ethanol in water along the side of the tube. Do not mix by pipetting. Centrifuge at >12,000 × g at 4 °C for 10 min.
b. 12. Carefully remove the supernatant as described in Step 10 and leave the tube to dry on the benchtop at room temperature for 5-10 min or until residual ethanol is no longer visible.
c. 13. Dissolve the RNA pellet in 50 µL of RNase-free water; mix by pipetting. Measure the concentration of RNA by using its absorbance at 260 nm using a NanoDrop spectrophotometer. Note that the A260/A280 ratio should be >1.9.

***Note:*** An aliquot (>200 ng) of total RNA should be saved as input to be compared with the pulled-down fractions. Steps 14-25 should be skipped, picking up the protocol again at Step 26 (Library Preparation and Sequencing).

**Pause point**: RNA samples can be stored at −20 °C for 1 week or at −80 °C for a month. For samples stored longer than these periods, the integrity of RNA should be assessed by using Bioanalyzer or gel electrophoresis prior to proceeding to the next steps.

Pull-down of cross-linked RNA fragments

### Timing: 8 h

The cross-linked and randomly fragmented RNA is captured onto azide-functionalized agarose beads by CuAAC. The unbound RNA fragments are washed away, and the captured RNA is released from the beads for downstream analysis. The following steps provide the amount of each material used for one sample containing 10 µg RNA. All centrifuge steps are performed by using a benchtop mini centrifuge (2000 × g) in this section unless otherwise noted.

a. 14. Transfer 100 µL of azide-disulfide agarose beads (Vector Laboratories, CCT- 1238) to a clean 1.6 mL microcentrifuge tube. The loading of these beads is 5- 20 µmol of azide groups per mL resin. Thus, 100 µL is sufficient to react with 0.5-2 µmol of alkyne groups from the chemical probe. [Note: following the procedure above, 0.2 mmol of probe was added per sample (20 mM × 10 mL).] Centrifuge for 30 s and remove the supernatant by pipet.

***Note***: The agarose beads are stored as a slurry and need to be gently mixed up before transferring. A P1000 tip is recommended for transferring to avoid clogging the tip. If needed, the stock agarose beads can be first diluted by adding 1 volume of 20% (v/v) ethanol in water to make pipetting easier.

a. 15. Add 200 µL 1× HEPES buffer (25 mM HEPES, pH 7.0) and invert the tube three times. Centrifuge for 30 s and remove the supernatant with a pipet.
b. 16. Add 10 µg of fragmented RNA to the beads in total volume of 50 µL, adjusting the volume with Nanopure water.
c. 17. In a separate tube, mix the following components for the click reaction:
d. 15 µL of 10 mM CuSO4
e. 15 µL of 50 mM THPTA
f. 15 µL of 250 mM sodium ascorbate. Although not adjusted, the pH should be ∼ 7.5.
g. 18. Add the click reaction mixture from Step 17 to the sample and incubate at 37 °C for 1 h while rotating or shaking (X rpm).
h. 19. Centrifuge for 30 s and remove the supernatant by pipet. Add 400 µL of High Salt Wash Buffer (10 mM Tris-HCl, 4 M NaCl, 1 mM EDTA, 0.1% Tween-20, pH 7.0) and invert the tube three times.
i. 20. Repeat Step 19 for a total of five washes. Remove the supernatant by pipet after the last wash.

***Note***: A small amount of residual liquid (<20 µL) is acceptable during the washes and to carry forward to Step #21 in order to minimize loss of beads. Spin filters may also be used to avoid bead loss.

a. 21. In a separate tube, mix the following components for cleaving the disulfide bond connecting the agarose beads and the “clicked” RNA. Bubbling is expected when the reagents are mixed together.
b. 50 µL of 200 mM TCEP in water
c. 50 µL of 100 mM K2CO3. The pH of this solution, although not adjusted, should be ∼11.
d. 22. Once bubbling has stopped, add the pre-mixed solution to the sample and incubate at 37 °C for 1 h while rotating or shaking (500 rpm).
e. 23. Add 50 µL of 200 mM iodoacetamide in water (light-sensitive) to the sample and incubate at 37 °C for 30 min while rotating or shaking (500 rpm) in the dark.

***Note***: Iodoacetamide is light-sensitive and should be stored at – 20 °C in the dark.

a. 24. Centrifuge for 30 s and carefully transfer the supernatant (now containing released RNA) with a pipet to a clean 1.6 mL microcentrifuge tube. A small amount of beads (<20 µL) carried over in the supernatant is acceptable, as they will be removed in the final clean-up step.
b. 25. Purify the RNA fragments by using RNA CleanXP beads (Beckman Coulter, A66514) following the manufacturer’s guide:
c. Add 270 µL of RNA CleanXP beads to the sample and mix well by pipetting up and down at least ten times. The resulting solution should be homogenous; if it is not continue to pipet until a homogenous solution is achieved. Incubate at room temperature for 5 min. As noted in the manufacturer’s protocol, addition of isopropanol can improve recovery of the RNA, although this is typically for short RNAs (<50 nt).
d. Place the tube onto the Agencourt SPRIstand (Beckman Coulter) magnetic rack for 5 min. Ensure that the beads are completely separated from the solution before proceeding to the next step.
e. Carefully remove the supernatant with a pipet without disturbing the beads.
f. Add 800 µL of 70% (v/v) ethanol to the beads. Make sure that all beads are submerged by the liquid. Per the manufacturer’s note, mixing the samples can reduce yield and is not recommended. Incubate at room temperature for 1 min. Carefully remove the supernatant with a pipet.
g. Repeat Step d for a total of three washes.
h. Dry the beads while the tubes are still in the magnetic rack by leaving the lids open to air for 5 – 10 min until no residual ethanol is visible.

**CRITICAL**: Incubating for longer than 10 min will over-dry the beads and cause poor RNA yield.

a. g. Add 50 µL of RNase-free water to elute the RNA from beads and pipet up and down gently a few times to mix while leaving tubes in the magnetic rack. Ensure all beads are rinsed by the water.
b. h. Incubate at room temperature for 5 min on the magnetic rack to separate beads from the solution, and then carefully transfer the solution to a clean tube. The expected RNA concentration is 10 – 50 ng/µL, as determined by measuring the absorbance by using Nanodrop UV spectrophotometer.

**Pause point**: RNA samples can be stored at −20 °C for 1 week or at −80 °C for a month. For samples stored longer than these periods, the integrity of RNA should be assessed by using a Bioanalyzer or gel electrophoresis prior to proceeding to the next steps.

***Note:*** The expected yield is 3 – 20%, depending on the selectivity of the chemical probe and abundance of the target RNA. If the yield is below 3%, increasing the amount of input RNA may provide sufficient RNA after pull-down for the steps below. However, a positive control (for example, the F1 compound reported here^1^ is expected to obtain

∼5% yield) should be used to confirm that the observed low yield is solely attributed to the nature of RNA-small molecule interactions, rather than technical problems. The integrity of the RNA after pull-down can be analyzed by gel electrophoresis or Bioanalyzer, and the expected average length is around 150 nt due to the fragmentation completed in Step #8. If the average length is below 50 nt, it is recommended to repeat the experiment as RNAs of this shortened length can reduce the number of mapped read counts in RNA-seq experiment.

Library preparation and sequencing

### Timing: 2 days

The steps below follow a standard pair-end RNA-seq workflow, which many institutional cores and companies can provide. Here, we describe depletion of rRNA from the samples, followed by library preparation using a NEBNext Ultra II Directional RNA Kit. All kits are commercially available. In our previously published work, libraries were analyzed using a NextSeq 500 v2.5 flow cell and sequenced with 2 × 40 bp paired-end protocol using a NextSeq sequencer. Longer reads (2 × 80 bp) are recommended, however, as they can improve the percentage of mapped reads in the downstream analysis.

***Note***: For each compound, both the RNA sample before the pull-down steps (see Note after Step 13) and after the pull-down (afforded after Step 25) must be sequenced to identify the enriched regions and reduce the false positives. A control probe lacking the RNA-binding module should also be analyzed in the same manner to identify background enrichment that is not driven by the (putative) RNA-binding module of the FFF or Chem-CLIP probe.

**CRITICAL**: All tubes and pipet tips must be free of any nucleases or oligonucleotide contamination.

a. 26. Quantify the concentration of pulled-down RNA, for example by using a Qubit

2.0 Fluorometer (Invitrogen), and measure the length distribution of the fragments, for example by using an Agilent 2100 Bioanalyzer RNA nanochip. We follow the standard manufacturer’s instructions for both quantification and length distribution (see Supplemental Information).

***Note***: The expected RNA concentration is 5 – 20 ng/µL. If RNA concentration is less than 5 ng/µL, the samples should be concentrated by a vacuum concentrator or by ethanol precipitation prior to proceeding to next steps. The distribution length of RNA samples should be centered around 150 – 200 nt (Figure 4). Additional fragmentation (Steps 8 - 13) can be performed again if the overall RNA length is too long. If the average length of RNA samples is below 50 nt, they are not suitable for proceeding to next steps (see Troubleshooting Problem 4). Note that RNA integrity numbers (RIN) are not suitable for assessing the integrity of these samples as they have been intentionally fragmented in Steps 8 – 13.

a. 27. Deplete rRNA from the sample by using NEBNext rRNA Depletion Module (E6310) per manufacturer’s recommended protocol as summarized below where the fragmentation steps have been omitted as they were already completed earlier in the protocol.

**Figure 4.**
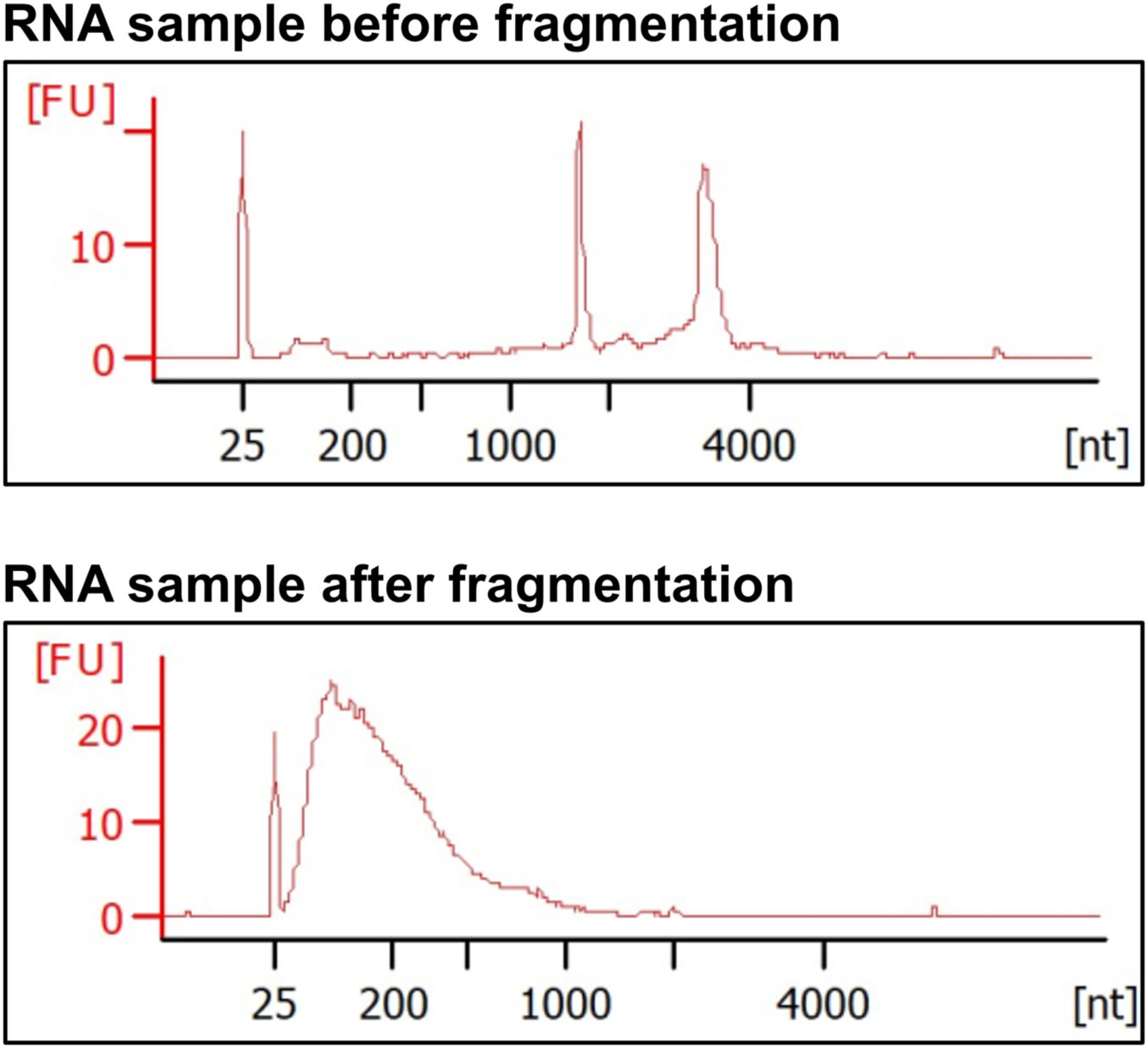
Example of RNA length distribution before and after random fragmentation to identify small molecule binding sites transcriptome-wide (Part II). Top, expected length distribution prior to fragmentation, as determined by Bioanalyzer analysis. Bottom, expected length distribution after fragmentation (∼150- 200 nt), as measured by Bioanalyzer (Steps #8 to 13 in Part II). Please refer to *Troubleshooting Part II Problem 4* if the length distribution is shorter or longer than expected.

a. Add 150 ng of RNA sample to a PCR tube containing 11 µL of nuclease-free water.

***Note***: The amount of input RNA can be lower if sufficient quantities were not obtained but should not be less than 50 ng.

i. b. Add 2 µL of NEBNext v2 rRNA Depletion Solution and 2 µL of NEBNext Probe Hybridization Buffer to the sample. Mix well by pipetting.
ii. c. Place the sample in a thermocycler with the following program (lid = 105

°C).

i. d. Briefly centrifuge the PCR tube to minimize loss of volume.
ii. e. Add 2 µL of RNase H Reaction Buffer, 2 µL of NEBNext Thermostable RNase H, and 1 µL of nuclease-free water to the sample. Mix well by pipetting at room temperature.
iii. f. Incubate the sample at 50 °C for 30 min.
iv. g. Add 5 µL of DNase I Reaction Buffer, 2.5 µL of NEBNext DNase I (RNase-free), and 22.5 µL of nuclease-free water to the sample. Mix well by pipetting.
v. h. Incubate the sample at 37 °C for 30 min.
vi. Briefly centrifuge the tube to minimize loss of volume.
vii. j. Purify the RNA by using RNAClean XP beads as described in Step 25 except using 90 µL of the beads in the first step and using 7 µL of nuclease-free water in the last step to elute RNA.

| Steps | Temperature | Time |
| --- | --- | --- |
| Initial Denaturation | 95 °C | 2 min |
| Hybridization | 0.1 °C/s | Ramp down to 22 °C |
| Hold | 22 °C | 5 min |

**Pause point**: RNA samples can be stored at −20 °C for 1 week or at −80 °C for a month. For samples stored longer than these periods, the integrity of RNA should be assessed by using a Bioanalyzer or gel electrophoresis prior to proceeding to the next steps.

i. 28. Prime the RNA by taking 5 µL of the sample from Step 27.
ii. Add 1 µL of Random Primers from the NEBNext Ultra II Directional RNA Kit to the sample and mix well by pipetting.
iii. Incubate the sample at 65 °C for 5 min and immediately transfer to ice.
iv. Centrifuge for 30 s by using a benchtop mini centrifuge (2000 × g).

***Note***: The fragmentation step from the manufacturer’s guide is omitted here as the RNA samples are already fragmented (Step 8).

i. 29. Keep the sample on ice and add the following components.
ii. 8 µL of NEBNext Strand Specificity Reagent
iii. 4 µL of NEBNext First Strand Synthesis Reaction Buffer
iv. 2 µL of NEBNext First Strand Synthesis Enzyme Mix
v. Mix the sample well by pipetting and centrifuge for 30 s by using a benchtop mini centrifuge (2000 × g). Place the samples in a thermocycler and run the following program:

i. 30. After centrifuging the samples (30 s in a benchtop mini centrifuge (2000 × g)), place the sample on ice and add the following components.
ii. 8 µL of NEBNext Second Strand Synthesis Reaction Buffer
iii. 4 µL of NEBNext Second Strand Synthesis Enzyme Mix
iv. 4 µL of nuclease-free water
v. 31. Mix well by pipetting and incubate at 16 °C for 1 h, preferably in a thermocycler. Centrifuge for 30 s by using a benchtop mini centrifuge (2000 × g).
vi. 32. Purify the double-strand cDNA by using NEBNext Sample Purification Beads.
vii. Add 144 µL of the beads to the sample and mix well by pipetting.
viii. Incubate at room temperature for 5 min.
ix. Place the tube on the magnetic rack to separate the beads. Carefully remove the supernatant with a pipet, without disturbing the beads.
x. Add 200 µL of 80% (v/v) ethanol and incubate at room temperature for 1 min. Remove the supernatant and repeat the wash once more (two total washes).
xi. Dry the beads by placing on the benchtops with the lids open for 5 min.
xii. Add 53 µL of the 0.1× TE Buffer (provided by the kit) and incubate at room temperature for 2 min. Ensure all beads are covered by the buffer.
xiii. Carefully transfer the supernatant containing the purified cDNA to a clean microcentrifuge tube.

| Steps | Temperature | Time |
| --- | --- | --- |
| 1 | 25 °C | 10 min |
| 2 | 42 °C | 15 min |
| 3 | 70 °C | 15 min |
| Hold | 4 °C | 5 min |

**Pause point**: Purified cDNA can be stored at −20 °C for 1 week or at −80 °C for 1 month. If longer storage period is needed, it is recommended to check the integrity of cDNA by using a Bioanalyzer or gel electrophoresis prior to proceeding to the next steps.

i. 33. Repair both ends of cDNA by adding the following components.
ii. 7 µL of NEBNext Ultra II End Prep Reaction Buffer
iii. 3 µL of NEBNext Ultra II End Prep Enzyme Mix
iv. 34. Mix well by pipetting and place in a thermocycler to run the following program.

i. 35. the samples from the centrifuge and place on ice. Add the following components to the sample.
ii. 5 µL of NEBNext Illumina Adaptor
iii. 1 µL of NEBNext Ligation Enhancer
iv. 30 µL of NEBNext Ultra II Ligation Master Mix
v. 36. Mix well by pipetting and incubate at 20 °C in a thermocycler for 15 min.

| Steps | Temperature | Time |
| --- | --- | --- |
| 1 | 20 °C | 30 min |
| 2 | 65 °C | 35 min |
| Hold | 4 °C | 5 min |

***Note***: The solution is viscous at this step and thorough mixing is required to ensure reaction efficiency.

i. 37. Add 3 µL of USER Enzyme to the sample. Mix well by pipetting and incubate at 37 °C for 15 min.
ii. 38. Purify the ligated cDNA as described in Step 33 except using 70 µL of beads in the first step and using 22 µL of 0.1× TE Buffer in the last step of elution.

**Pause point**: Purified cDNA can be stored at −20 °C for 1 week or at −80 °C for 1 month. If longer storage period was used, it is recommended to check the integrity of cDNA by using a Bioanalyzer or gel electrophoresis prior to proceeding to the next steps.

i. 39. Add the following component to a PCR tube.
ii. 22 µL of the purified cDNA from Step 39.
iii. 25 µL of NEBNext Ultra II Q5 Master Mix
iv. 5 µL NEBNext Primer Mix
v. 40. Place the sample onto a thermocycler and execute the following program.

| Steps | Temperature | Time | Cycles |
| --- | --- | --- | --- |
| Initial Denaturation | 98 °C | 30 sec | 1 |
| Denaturation | 98 °C | 10 sec | 12-13 cycles |
| Annealing/Extension | 65 °C | 75 sec |  |
| Final extension | 65 °C | 5 min | 1 |
| Hold | 4 °C | 5 min |  |

**CRITICAL**: The number of PCR cycles is dependent on the input RNA (150 ng for this protocol). Overamplification can significantly affect sequencing results.

**Pause point**: The PCR product can be stored at −20 °C for 1 week or at −80 °C for 1 month. If longer storage period was used, it is recommended to check the integrity of cDNA by using a Bioanalyzer or gel electrophoresis prior to proceeding to the next steps.

i. 41. Purify the PCR product as described in Step 33, except with the following changes:
ii. Use 45 µL of beads in the first step.
iii. Use 23 µL of 0.1× TE Buffer in the last step for elution.

***Note***: If significant primer dimer (∼80 bp) or adaptor dimer peaks (∼150 bp) are observed, the PCR product must be repurified by size-selection beads or by denaturing polyacrylamide gel electrophoresis.

i. 42. Analyze the length of the purified PCR product using an Agilent 2100 Bioanalyzer, which should show a peak distribution centered at ∼300 bp.
ii. 43. Load the final library into a NextSeq 500 v2.5 flow cell and sequence with the Illumina pair-end 80 bp program.

RNA-seq data analysis

## Timing: 1 – 2 days

The RNA-seq data are aligned to the human genome by using STAR,^13^ and enriched regions are identified by using Genrich (available from github.com/jsh58/Genrich). Enrichment is calculated by using the bamCompare.^14^ The sequencing track can be visualized by using the IGV browser.^15^

RNA-seq analysis is mainly performed by using high performance computer (HPC) while some steps can be performed locally on personal laptops. Note that this protocol is written for Windows users and some commands may be different for Linux or Mac platforms. A basic overview of RNA-seq analysis can be found from rnaseq.uoregon.edu, and a virtual open course for learning RNA-seq analysis can be found from diytranscriptomics.com.

***Note***: No unique codes are used for this analysis. Below are standard scripts adapted from the user manual for each algorithm. It is strongly recommended to read the published manual (Supplemental Information) carefully to fully understand the algorithm as well as troubleshooting. If error messages occur during the RNA-seq analysis, please refer to the user manual for each package (Supplemental Information) as well as online forums such as Biostars (www.biostars.org/t/Forum) and SEQanswers (www.seqanswers.com/forum/applications-forums/rna-sequencing) for troubleshooting.

i. 44. Select the tool to align RNA-seq data to the reference genome. A number of different alignment tools are available for this purpose, including STAR (github.com/alexdobin/STAR/tree/master), bwa (github.com/lh3/bwa), HISAT2 (daehwankimlab.github.io/hisat2), salmon (combine-

lab.github.io/salmon/about), and Kallisto (github.com/pachterlab/kallisto). A comprehensive review and comparison of these different alignment tools can be found here.^16^ For this protocol, STAR is used as the alignment tool.

***Note:*** STAR was chosen over other tools due to: (1) its overall superior performance in alignment accuracy; (2) tunable parameters for different experimental settings; (3) regular maintenance by the developer; and (4) clear user manual and guidance with abundant online resources for tips and troubleshooting. The main disadvantage of STAR is the high cost of memory and computational resources, which can be accommodated by using HPC. While possible, it is not recommended to install and run STAR on a local PC.

1. 45. Create an index file. Prior to the alignment, STAR requires a map between the transcriptome and the genome of any organism of interest, a file that is known as “index file”. The index file is constructed by STAR by using publicly available transcriptome and genome database. The most updated versions of the genome files can be downloaded from www.gencodegenes.org.
2. For each organism, two files need to be downloaded, one ending with .fa (the sequences) and the other ending with .gtf (the annotation of sequences). For example, for human (*Homo sapiens*), these two files are:
3. GRCh38.primary_assembly.genome.fa
4. gencode.v38.primary_assembly.annotation.gtf

***Note:*** *GRCh38* and *v38* refer to the version number and must be the same number for the two files. While these files refer to the most used human genome for RNA-seq analysis, two recently assembled human genome can also be used: (1) the Telomere-to-Telomere (T2T) genome,^17^ which provides a more comprehensive annotation of the human genome, especially for the difficult-to-sequence regions; and (2) the Human Pangenome,^18^ which provides annotation from a cohort of genetically diverse individuals to account for genetic diversity in human populations.

1. b. Unzip the downloaded files by using the following command.
2. c. Generate the index by running the following command. Please allow a minimum of 40 GB RAM to perform this step, which can take 30 – 120 minutes depending on the computing power and the size of the genome files. The parameters used in the last two lines are recommended by the STAR manual. In particular “sjbdOverhang 99” specifies the length of the genomic sequence around the annotated junction to be used in constructing the splice junctions database. In principle, this length should be equal to the length of the reads minus 1. However, the developer recommends using 99 for all standard Illumina Nextseq files (whether 2 × 40 or 2 × 80 bp sequencing set-up). This parameter may need to be adjusted if long-read sequencing (>100 bp) was used, in which case please refer to the developer’s website (github.com/alexdobin/STAR) for advice. The “genome ChrBinNbits 18” specifies the size of the bins for genome storage. The default value of 18 is suitable for most species including humans and mice. If genome files with a large number of references are used (*i.e.*, a large number of chromosomes and annotations), reducing this value may help reduce RAM consumption. For more information about these parameters, please refer to the STAR manual (Supplemental Information).

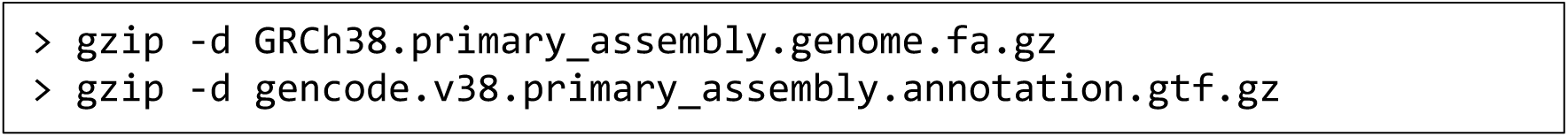

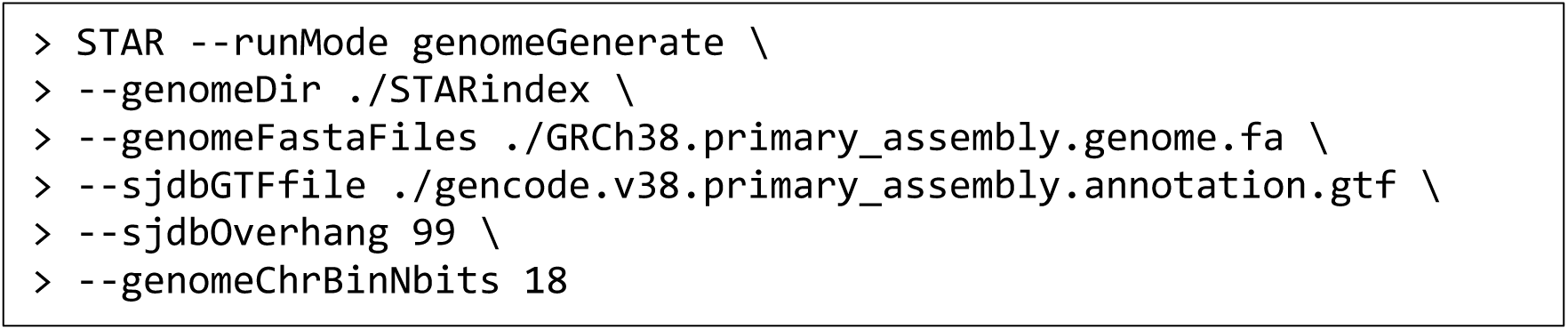

***Note:*** Step #45 only needs to be performed once and the index file can be re-used for additional analyses. However, a new index file needs to be created by repeating Step #45 if any of the following scenarios applies: (1) changing the species; (2) changing the versions of the public genomic files; (3) updating the versions of the STAR package.

1. 46. Align the fastq files to the human genome.

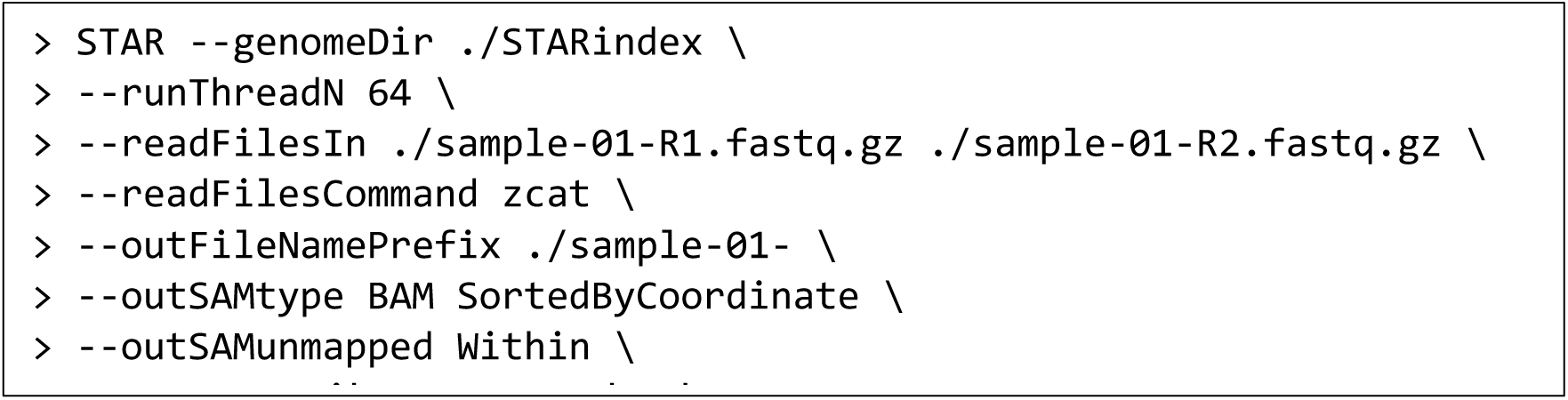

Brief explanations of each parameter are provided below and a comprehensive explanation can be found at STAR manual (Supplemental information):

*--genomeDir* is where the index file is located.

*--runThreadN 64* means using 64 threads in parallel. The larger this number is, the faster computation will be, however the maximum number of threads depends on the HPC capacity.

*--readFilesIn* is where the input files are located. For paired-end sequencing, two raw sequencing files are obtained and typically annotated by R1 (read 1) and R2 (read 2).

*--readFilesCommand zcat* means the input files are compressed. (Note that these fastq files end with .gz indicating they are compressed to save space.)

*--outFileNamePrefix* is the prefix of the output file after alignment.

*--outSAMtype* is the type of output files.

***Note:*** The .bam file is the main output file containing all aligned sequences to the genome of interest.

The Log.final.out file contains summary information for a quality control check. This file can be opened as a .txt file either on HPC or the local PC. A few variables in this table should be checked for quality control including the following:

- *Number of input reads:* This should match the estimated read depth from the sequencing core, usually ∼20 million per sample for total RNA-seq.
- *Average input of read length:* This should match the sequencing kit used by the core.
- *Uniquely mapped reads%:* This percentage shows how many reads are uniquely mapped to the genome (*i.e.*, there is no ambiguity in mapping these reads to a gene). A higher percentage of uniquely mapped reads indicates higher data quality. The recommended percentage is >80%, and an acceptable minimum is 60%, although there is no consensus in the field regarding these values. There are a few factors that can affect this value, including RNA integrity, the library preparation, and the read length.

***Note:*** This process is also known as “Peak Calling”, and the algorithms used in this section are adapted from cross-linking and immunoprecipitation (CLIP) experiments used to study RNA-binding proteins. A number of packages can be used for this purpose, including CLIP Tool Kit (zhanglab.c2b2.columbia.edu/index.php/CTK_Documentation) and Model-based Analysis of ChIP-Seq (MACS).^19^ In this protocol, Genrich (github.com/jsh58/Genrich) was selected as it is widely used with clear documentation. The workflow of Genrich and comprehensive explanations can be found at its webpage (github.com/jsh58/Genrich).

1. 47. Identify the regions enriched by using Genrich. The aligned bam files are parsed by another algorithm to look for enriched regions based on statistical analysis of read counts from samples before and after the Chem-CLIP process. Here “sample-01” refers to the RNA after pull-down, and “sample-02” to the input RNA prior to pull-down. It is recommended to apply manual filtering by using MS Excel to eliminate low-confidence peaks, *i.e.*, by applying a minimum read count of 5 and a minimum fold of enrichment of 1.5, although different cut-offs can be chosen to increase or decrease stringency.

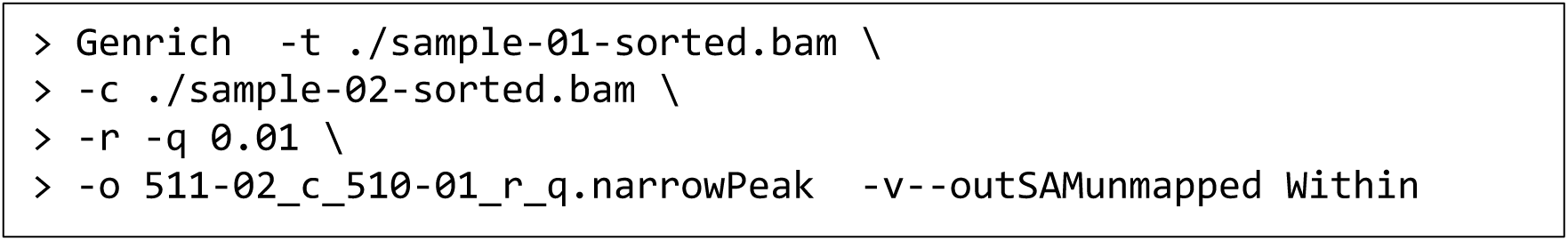

***Note:*** Genrich has the following limitations:

- The identified regions of enrichment (peaks) are based on mathematical models, so the output coordinates may not exactly match the shape of peaks. To address this issue, see the next section on visualization.
- Genes with low read coverage are prone to be identified as peaks due to large variance of read counts. To address this issue, a minimum read of 5 is applied to manually remove lowly expressed targets from further analysis.
- Regions near the ends of the transcript (5’ and 3’ untranslated regions (UTRs)) may be prone to potentially higher false positive rates due to less sequencing coverage towards the ends of a transcript. This cannot be easily resolved as it is intrinsic to the principles of NGS library preparation; thus care should be taken when inspecting peaks near 5’ or 3’ UTR.
- Lowly expressed RNA targets may not receive sufficient reads from RNA-seq and therefore are more prone to false negatives in this analysis.
- 48. Quantify the fold of enrichment by using bamCompare. A full explanation of bamCompare can be found at its website (deeptools.readthedocs.io/en/develop/content/tools/bamCompare). Briefly, it compares two files based on the number of mapped reads. The genome is partitioned into bins of equal size, then the number of reads found in each bin is counted per file, and a ratio between for the number of reads found in each bin between two files are reported. This tool accounts for the normalization of different read depth from different samples by using a statistical method reported previously.^20^

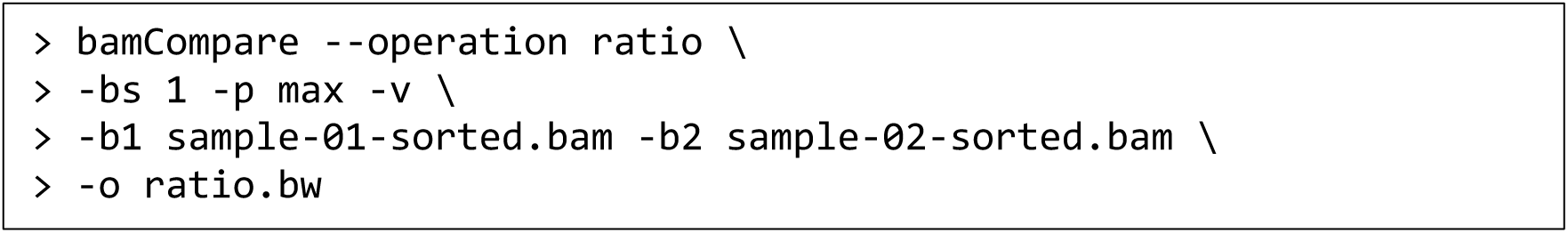

*-- operation ratio* calculates the ratio between these two files (*i.e.*, the bam file after vs. before pull-down).

-b1 is after pull-down while -b2 is input (before pull-down)

*-bs 1* means the bin size is set to 1 (single nucleotide). Increasing the bin size can save calculation time but will reduce resolution of the reads. (Note that this does not necessarily imply single nucleotide resolution of an RT stop that indicates the binding site.)

*-p max* means this task will be run utilizing the maximum number of available threads.

*-v* means the real-time progress will be shown on the user’s screen.

*-o* refers to the output file location and name (default is at current location).

The output file is .bw that contains the ratio of reads after vs. before pull-down across the entire genome, which can be opened by importing into MS Excel or by IGV browser.

***Note:*** the following suggestions and considerations are recommended when running bamCompare.

- If only a specific region of the genome is of interest, use -r to specify this region and therefore save the calculation time. For example, –r chr10 to only calculate chromosome 10.
- To avoid mathematical errors when dividing over zero, any region that has zero read will be automatically assigned as 1 read count. This can lead to artificially high ratio for the regions with low or zero read coverage in the sample before pulldown. A minimum of 5 reads after pull-down in the regions of interest is recommended to be considered as potentially enriched.
- The fold of enrichment can be reported by either the peak value (the point of highest ratio) within the region of enrichment, or the average value across the region of enrichment. In many cases, these two values are similar. However, one should be consistent with this choice when analyzing different targets.
- For an identified region with enrichment by the compound, it is important to check the same region from the control probe samples. A minimum of 3-fold enrichment difference between the compound and the control probe is recommended to consider a target as being *bona fide* target of the compound rather than from background enrichment due to non-specific reaction of the diazirine.
- 49. Identify RT-stop (binding) sites. Visualize the bam and bw files by using IGV browser. Comprehensive instructions for using the IGV browser can be found at its website (www.igv.org). Zoom in to the peaks identified in Step #48 and the mapped RT-stop site is the end of the peak (exact chromosomal coordinates are reported in the output files from Step #48).

***Note:*** Although Step #48 reports single-nucleotide coordinates based on statistical significance, the current experimental workflow does not necessarily provide single-nucleotide resolution of mapped RT-stops. One should also be cautious of identified RT-stop sites near 5’ or 3’ ends of the transcript, as these regions may be prone to false positives due to less sequencing coverage. The following two scenarios should be considered as red flags for an identified RT-stop as potential false positives: (1) the control probe group showing exactly the same (or nearly the same) RT-stop site; (2) the RT-stop site is at the exon-intron junction, where the read coverage naturally drops due to the relative high abundance of expressed exons to introns. Note that not all enriched RNA targets can be precisely mapped with the small molecule binding sites, potentially due to the flexible structures formed near the enriched region that lacks a defined binding site.

1. 50. Model the structure of the RNA around the mapped binding sites. In this published study,^1^ a computational pipeline known as ScanFold^21^ was used to model the RNA structures around the small molecule binding sites. Compared to other MFE-based RNA folding algorithms, ScanFold integrates evolutionary selection to identify regions with high probability of biological function. A detailed explanation and usage of ScanFold is available from its website: mosslabtools.bb.iastate.edu. For this published study,^1^ a default window size of 120 nt is used, which is recommended by the developers of ScanFold.

Expected outcomes

**Part I:** This protocol describes a method to measure RNA target occupancy by small molecules *in vitro* by capturing a covalent cross-link induced by UV irradiation. The percentage of RNA pulled down can vary widely, depending on target and probe interactions (affinity, on- and off-rates, etc.), as well as probe concentrations. When designing a Chem-CLIP probe based on a small molecule of interest, the attachment of the photoaffinity label can impact target engagement and, if possible, it is recommended to confirm that such modification is tolerated, e.g. by maintaining biological activity. It should be noted that even very similar Chem-CLIP probes can have very different labeling intensities.^22^ Conversely, the attachment point of the cross- link and enrichment handle of a hit compound from an FFF library screening, presents a convenient opportunity to create a modularly assembled small molecule that binds two sites simultaneously^23^ or develop RNA degraders.^24^

The background of pull-down observed in the vehicle (DMSO-treated) sample is typically ∼1% and should not exceed 5%.

**Part II:** This protocol provides a method to map the binding sites of small molecules to RNA within the human transcriptome in an intact biological system. The total RNA yield from one 100 mm diameter dish is ∼15 µg for MDA-MB-231 cells, however the yield can vary dramatically depending on cell type/line. The yield after pull-down is ∼5% and depends on the selectivity of the compound; that is, a promiscuous RNA binder is expected to afford a higher yield after pull-down than a selective small molecule.

At various points in the protocol, the fragmented RNA, cDNA, and PCR product are purified by either ethanol precipitation or magnetic beads, and the yield is expected to be >90%. Figure 4 shows the expected length distribution of RNA sample before and after the random fragmentation (Step 13).

The expected number of reads from Nextseq 500 is around 20 million per sample, although deeper sequencing could be used for lowly expressed RNA targets. In Step 45, the percentage of uniquely mapped reads is expected to be >50%. Figure 5 shows an example of the sequencing tracks of an identified enriched target as well as the mapped binding site. Note that skyline plots in panel A and in the first two rows (before and after pull-down) of panel B are not normalized for read depth, while the skyline plot on the last row (ratio of enrichment) of panel B is normalized for the read depth of each sample. Therefore, the shape of peaks shown on the skyline plot on last row (ratio of enrichment) of panel B is different from simply dividing the middle row (after pull-down) by the top row (before pull-down).

**Figure 5.**
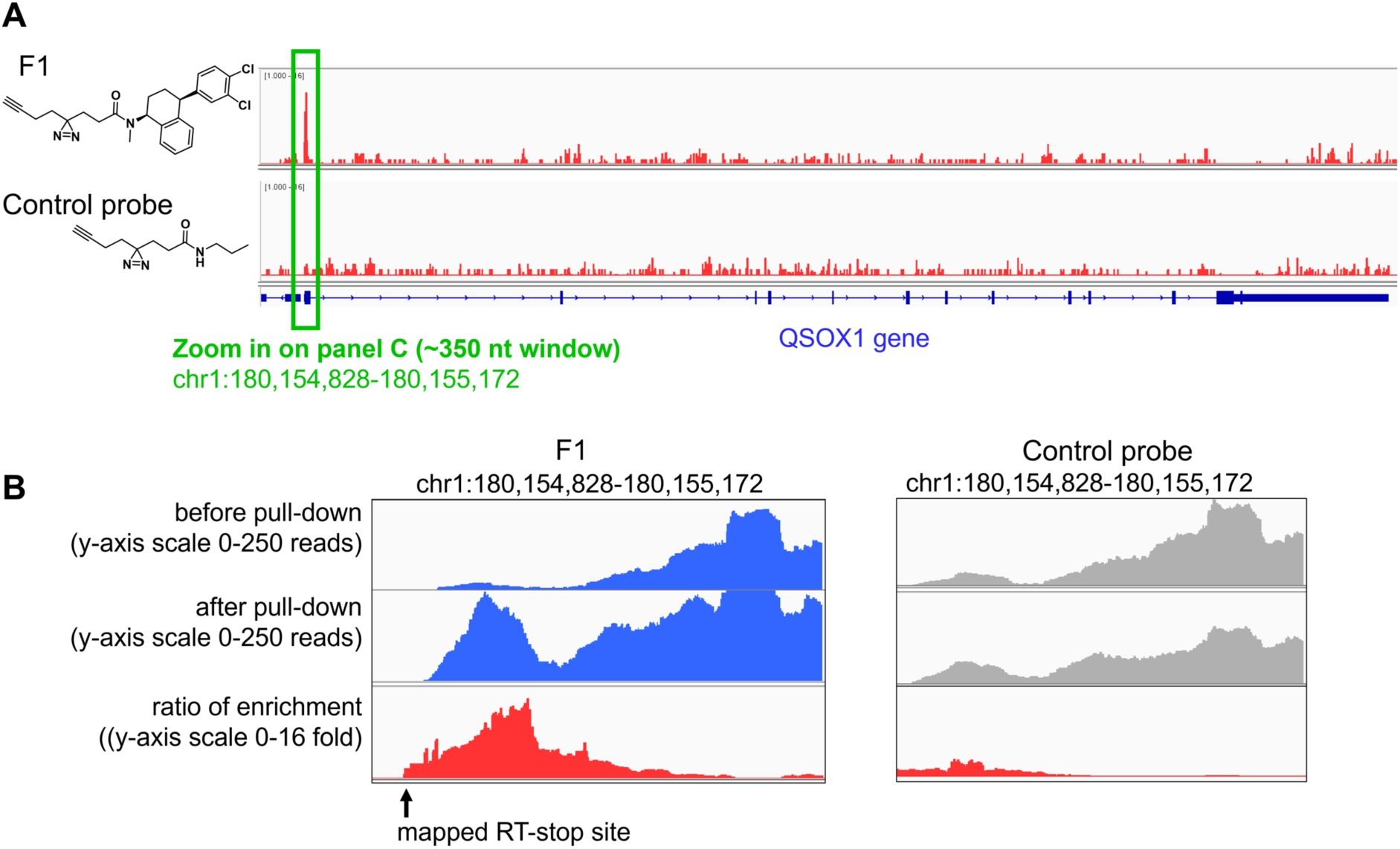
The example skyline plot of the enriched target and the mapped binding site (Part II). Example of F1 enriching *QSOX1-a* mRNA in MDA-MB-231 cells, as reported previously.^1^ (A) Structures of the small molecule probe F1 and the control probe lacking the RNA-binding module and the corresponding skyline plot of the QSOX1 gene visualized by IGV browser for the output from Step #49. A peak of enrichment was identified (green box) in the F1-treated group but not in the control group. (B) Zoom- in to the region of enrichment by using IGV browser. The mapped RT-stop site is reported from step #47 by Genrich based on statistical significance. Note that skyline plots in panel A and shown in the first two rows (before and after pull-down) of panel B are not normalized for read depth, while the skyline plot on the last row (ratio of enrichment) of panel B is normalized for the read depth of each sample. Therefore, the shape of peaks shown on the skyline plot on last row (ratio of enrichment) of panel B is different from simply dividing the middle row (after pull-down) by the top row (before pull-down).

Limitations

**PART I:** Depending on both target and probe, the cross-linking efficiency might be very low, as the diazirine group may be quenched by water molecules in the solution or undergo other side reactions.^25, 26^ Alternative cross-linking groups such as chlorambucil^5^ should be considered, although its head-to-head comparisons with diazirine has not yet been performed. Nitrogen mustards spontaneously form a reactive aziridinium ion in the aqueous buffer,^27^ whereas diazirines produce short-lived carbenes and diazo species upon UV irradiation.^28^ While N7-G is generally considered the most nucleophilic position in the nucleobases, and has been reported as the predominant site to react with nitrogen mustards,^29^ diazirines have been shown to be able to react with all nucleobases,^30^ and are generally able to insert into C-H bonds.^26^ Therefore, these photoaffinity labels are expected to be less limited in their reaction partners, while reactivity biases might still exist as has been observed for proteins.^25^ Further points of consideration for the use of nitrogen mustards are reaction with intrinsically more reactive RNA sites independent from small molecule binding (for diazirines rapid quenching should mitigate this), potentially longer incubation times required for reaction to reach completion, and potentially larger contributions to target interactions by the crosslinking module (overall size, stacking and charge interactions). Furthermore, if validation of target engagement in live cells is desired, diazirines are to be expected to have a better toxicity profile than nitrogen mustards that can potentially cause DNA damage.

It is also of note that, the interactions observed in this limited *in vitro* setting might not reflect conditions in live cells, due to different RNA folding and additional interaction partners such as RNA-binding proteins or other RNAs. Nevertheless, the experiment is straightforward and can serve as a screening platform for the streamlined identification of lead compounds and enables the validation of target engagement.

**PART II:** This protocol is designed to define the cellular molecular fingerprints of small molecules within the human transcriptome. However, a few limitations should be considered when applying this method. One limitation is the cross-linking efficiency of the diazirine module which may not be sufficient to capture all the bound targets, especially ones with low abundance or short half-lives. Further, the current experimental workflow removes short RNA such as microRNAs and tRNAs in the library preparation step, and thus these short RNA targets will not be assessed in the downstream analysis. Previous studies also demonstrated that the linker length between the diazirine module and the RNA-binding module can significantly affect which targets are cross-linked.^22^ Another limitation is that not all enriched RNA targets can be precisely mapped with the small molecule binding sites. This is potentially caused by the flexible structures formed near the enriched region that lacks a defined binding site. RNA-seq analysis is a rapidly evolving field and discussion on its limitations can be found elsewhere.^31^

### Troubleshooting

Part I: in vitro Chem-CLIP with radiolabeled RNA Problem 1:

No pull-down observed.

Potential solution:

Attempt alternative buffer systems, ensuring that the RNA construct is folded properly under the experimental conditions. Alternative probes might result in more efficient cross-linking.^22^

Problem 2:

Pull-down is observed in control samples, such as DMSO, “no UV”, or “no click”.

Potential solution:

Ensure that no ambient light is responsible for photoactivation. While the described washing conditions already are stringent, different reaction buffers and/or wash buffers could help reduce non-specific pull-down. If optimizing washing conditions, it is recommended to collect each individual wash in a separate scintillation vial.

Part II: Transcriptome-wide mapping in live cells Problem 1:

Low RNA yield after extraction from the cells after UV cross-linking. Potential solution:

Some cells may easily detach during cross-linking, which can be confirmed by using a microscope. In this case, do not discard the 1× DPBS in the dish. Gently scrape the dish to detach all the cells into the 1× DPBS and then transfer the buffer to a clean conical vial. Centrifuge at 160 × g for 5 min and then remove the supernatant with a serological pipet. Proceed to RNA extraction as described in Step 6a. It should also be noted that toxicity of the compound could also lead to low cell density and therefore low RNA yield. To avoid toxicity, consider reducing the compound concentration and/or reducing the treatment time. Finally, consider increasing the number of dishes and combine multiple dishes as one biological replicate to obtain sufficient RNA for further analysis.

Problem 2:

Low RNA yield after pull-down by azide-functionalized beads. Potential solution:

Ensure that the azide-functionalized agarose beads are well mixed before pipetting (P1000 pipet is recommended to avoid clogging on the tip). Avoid mechanical stirring or vortexing to avoid shattering or shearing the beads. Ensure the beads are stored at 4 °C and not frozen.

Problem 3:

Low RNA quality with partially degraded RNA before random fragmentation. Potential solution:

Ensure the working environment is free of RNases. Keep RNA samples on ice unless actively working on the sample.

Problem 4:

The average length of RNA samples is too short (<50 nt) or too long (>200 nt) at Step #26.

Potential solution:

For RNA samples that are too short, repeat the experiment by reducing the fragmentation time at Step #8. Ensure the working environment is free of RNases. Keep RNA samples on ice unless actively working on the sample. For RNA samples that are too long, additional fragmentation can be performed at Step #26. If repeating the experiment, increase the fragmentation time at Step #8.

Problem 5:

Low percentage of reads mapped after Step #45. Potential solution:

Purify the final library by denaturing polyacrylamide gel electrophoresis to remove primers and adaptor dimers.

## Resource availability

### Lead contact

Further information and requests for resources and reagents should be directed to and will be fulfilled by the lead contact, Matthew D. Disney.

### Materials availability

All materials are available upon reasonable request to the lead contact.

### Data and code availability

No unique codes were used in this study. The full dataset, which was previously published,^1^ is deposited on Mendeley Data (DOI: 10.17632/56r9zmjps2.1).

## Acknowledgments

This work was supported by the National Institutes of Health (R01 CA249180A and R35 NS116846 to M.D.D.) and the Department of Defense (W81XWH-20-1-0727 and HT94252310336).

## Author contributions

M.D.D. conceived the idea and directed the study. Y. T., P.R.A.Z., X.Y, X.S. performed the experiments and data analysis. J. L. C-D. edited the manuscript with the help from all authors.

