## Supplementary File 1 for "Transcriptome-wide mapping of small-molecule RNA-binding sites in live cells"

### RNA 6000 Nano Kit for 2100 Bioanalyzer Systems

### Quick Guide

The complete *RNA 6000 Nano Kit for 2100 Bioanalyzer Systems Kit Guide* can be found in the online help of the Agilent 2100 Expert software.

#### Kit Components

| **Agilent RNA 6000 Nano Kit (5067-1511)** | |
|---|---|
| **Agilent RNA 6000 Nano Chips** | **Agilent RNA 6000 Nano Reagents (5067-1512) & Supplies** |
| 25 RNA Nano Chips | ● (yellow) RNA 6000 NanoLadder (1 vial, 5067-1529) |
| 2 Electrode Cleaners | ● (blue) RNA 6000 Nano Dye Concentrate (1 vial) |
| | ● (green) RNA 6000 Nano Marker (2 vials) |
| **Syringe Kit** | ● (red) RNA 6000 Nano Gel Matrix (2 vials) |
| 1 Syringe | 4 Spin Filters (5185-5990) |

| **Tubes for Gel-Dye Mix** |
|---|
| 30 Safe-Lock Eppendorf Tubes PCR clean (DNase/RNase free) for gel-dye mix |

#### For Research Use Only

Not for use in Diagnostic Procedures.

#### Assay Principles

Agilent RNA kits for the 2100 Bioanalyzer system contain chips and reagents designed for sizing and analysis of RNA fragments. Each chip contains an interconnected set of microchannels that is used for separation of nucleic acid fragments based on their size as they are driven through it electrophoretically. This kit is designed for use with the 2100 Bioanalyzer system only.

#### Applications and Kits

Agilent RNA kits are designed for the analysis of total RNA (eukaryotic, prokaryotic, and plant) and mRNA samples. Available kits: Agilent RNA 6000 Nano kit (5067-1511), RNA 6000 Pico kit (5067-1513) and Small RNA kit (5067-1548)

#### Storage Conditions

- Freeze unopened RNA ladder at -28 – -15 °C (-18 – 5 °F). Prepared ladder aliquots need to be stored at -28 – -15 °C (-18 – 5 °F). Keep all other reagents and reagent mixes refrigerated at 2 – 8 °C (36 – 46 °F) when not in use to avoid poor results caused by reagent decomposition.

- Protect dye and dye mixtures from light. Remove light covers only when pipetting. Dye decomposes when exposed to light.
- Store the chips at room temperature.

#### Equipment Supplied with the Agilent 2100 Bioanalyzer System

- Chip priming station (5065-4401)
- IKA vortex mixer

#### Additional Material Required (Not Supplied)

- RNaseZAP® recommended for electrode decontamination and routine electrode cleaning
- RNase-free water recommended for routine electrode cleaning
- Pipettes (10 µL and 1000 µL) with compatible tips (RNase-free, no filter tips, no autoclaved tips)
- 0.5 mL and 1.5 mL microcentrifuge tubes (RNase-free)
- Microcentrifuge (>13000 g)
- Heating block or water bath for ladder/sample preparation

#### Sample preparation

For total RNA or mRNA analysis, the sample concentration must be within the specified range. If the concentration of your particular sample is above this range, dilute with RNase-free water.

#### Specifications

| Physical Specifications | | Analytical Specifications | | |
|---|---|---|---|---|
| | | | **Total RNA Assay** | **mRNA Assay** |
| Analysis time | 30 min | Quantitative range | 25 – 500 ng/µL | 25 – 250 ng/µL |
| Samples per chip | 12 | Qualitative range | 5 – 500 ng/µL | 5 – 250 ng/µL |
| Sample volume | 1 µL | Sensitivity (S/N>3) | 5 ng/µL in water | 25 ng/µL in water |
| Kit stability | 4 months | Quantitative precision (within a chip) | 10 % CV | 10 % CV |
| Kit size | 25 chips<br>12 samples/chip<br>= 300 samples/kit | Quantitative accuracy[1] | 20 % | 20 % |
| | | Maximum salt concentration in sample | 100 mM Tris<br>0.1 mM EDTA<br>or 125 mM NaCl<br>15 mM $MgCl_2$ | 100 mM Tris<br>0.1 mM EDTA<br>or 125 mM NaCl<br>15 mM $MgCl_2$ |

1 Determined analyzing the RNA ladder as sample

#### Setting up the Chip Priming Station

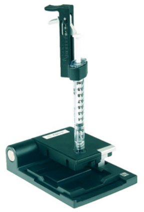

1 Replace the syringe:
   a Unscrew the old syringe from the lid of the chip priming station.
   b Release the old syringe from the clip. Discard the old syringe.
   c Remove the plastic cap of the new syringe and insert it into the clip.
   d Slide it into the hole of the luer lock adapter and screw it tightly to the chip priming station.

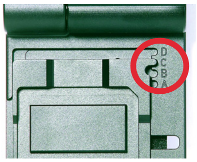

2 Adjust the base plate:
   a Open the chip priming station by pulling the latch.
   b Using a screwdriver, open the screw at the underside of the base plate.
   c Lift the base plate and insert it again in position C. Retighten the screw.

3 Adjust the syringe clip:
  a Release the lever of the clip and slide it up to the top position.

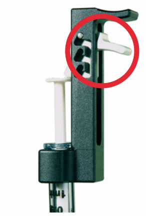

#### Essential Measurement Practices

- Handle and store all reagents according to the instructions on the label of the individual box.
- Avoid sources of dust or other contaminants. Foreign matter in reagents and samples or in the wells of the chip will interfere with assay results.
- Allow all reagents to equilibrate to room temperature for 30 min before use. Thaw samples on ice.
- Protect dye and dye mixtures from light. Remove light covers only when pipetting. The dye decomposes when exposed to light and this reduces the signal intensity.
- Always insert the pipette tip to the bottom of the well when dispensing the liquid. Placing the pipette at the edge of the well may lead to poor results.
- Always wear gloves when handling RNA and use RNase-free tips, microcentrifuge tubes and water.
- It is recommended to heat denature all RNA samples and RNA ladder before use for 2 min and 70 °C (once) and keep them on ice.
- Do not touch the 2100 Bioanalyzer instrument during analysis and never place it on a vibrating surface.
- Always vortex the dye concentrate for 10 s before preparing the gel-dye mix and spin down afterwards.
- Use a new syringe and electrode cleaners with each new kit.
- Use loaded chips within 5 min after preparation. Reagents might evaporate, leading to poor results.
- To prevent contamination (e.g. RNase), it is strongly recommended to use a dedicated electrode cartridge for RNA assays.
- Perform the RNase decontamination procedure for the electrodes daily before running any assays. Refer to the kit guide for details on electrode cleaning and decontamination.

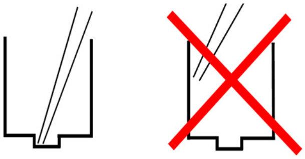

#### Agilent RNA 6000 Nano Assay Protocol

**WARNING**

**Handling Reagents**

**The dye can cause eye irritation. Because the dye binds to nucleic acids, it should be treated as a potential mutagen.**

**Kit components contain DMSO. DMSO is skin-permeable and can elevate the permeability of other substances through the skin.**

- ✓ **Follow the appropriate safety procedures and wear personal protective equipment including protective gloves and clothes as well as eye protection.**
- ✓ **Follow good laboratory practices when preparing and handling reagents and samples.**
- ✓ **Always use reagents with appropriate care.**
- ✓ **For more information, refer to the material safety data sheet (MSDS) on www.agilent.com.**

#### Preparing the RNA Ladder

1. Spin the ladder down and pipette in an RNase-free vial.
2. Heat denature the ladder for 2 min at 70 °C.
3. Immediately cool the vial on ice.
4. Prepare aliquots in recommended 0.5 mL RNase-free vials with the required amount for typical daily use.
5. Store aliquots at -28 – -15 °C (-18 – 5 °F). After initial heat denaturation, the frozen aliquots should not require repeated heat denaturation.
6. Before use, thaw ladder aliquots on ice (avoid extensive warming).

#### Preparing the Gel

1. Pipette 550 µL of RNA gel matrix (red ●) into a spin filter.
2. Centrifuge at 1500 g ± 20 % for 10 min at room temperature.
3. Aliquot 65 µL filtered gel into 0.5 mL RNase-free microcentrifuge tubes. Use filtered gel within 4 weeks. Store at 2 – 8 °C (36 – 46 °F).

#### Preparing the Gel-Dye Mix

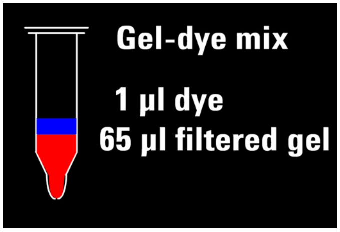


1. Allow the RNA dye concentrate (blue ●) to equilibrate to room temperature for 30 min.
2. Vortex RNA dye concentrate (blue ●) for 10 s, spin down and add 1 µL of dye into a 65 µL aliquot of filtered gel.
3. Vortex solution well. Spin tube at 13000 g for 10 min at room temperature. Use prepared gel-dye mix within one day.

#### Loading the Gel-Dye Mix

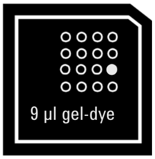


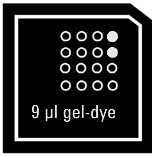


1. Put a new RNA chip on the chip priming station.
2. Pipette 9 µL of gel-dye mix in the well marked G.
3. Make sure that the plunger is positioned at 1 mL and then close the chip priming station.
4. Press plunger until it is held by the clip.
5. Wait for exactly 30 s then release clip.
6. Wait for 5 s. Slowly pull back plunger to 1 mL position.
7. Open the chip priming station and pipette 9 µL of gel-dye mix in the wells marked G.
8. Discard the remaining gel-dye mix.

#### Loading the Marker

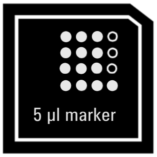


1. Pipette 5 µL of RNA marker (green ●) in all 12 sample wells and in the well marked (ladder).

#### Loading the Ladder and Samples

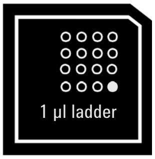


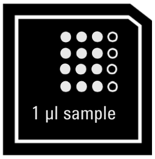


1. Pipette 1 µL of prepared ladder in well marked (ladder).
2. Pipette 1 µL of sample in each of the 12 sample wells. Pipette 1 µL of RNA marker (green ●) in each unused sample well.
3. Put the chip horizontally in the IKA vortexer and vortex for 1 min at 2400 rpm.
4. Run the chip in the 2100 Bioanalyzer instrument within 5 min.

#### Technical Support

Please visit our support web page **www.agilent.com/genomics/contactus** to find information on your local Contact Center.

#### Further Information

Visit the Agilent website. It offers useful information, support, and current developments about the products and technology: **www.agilent.com/en/product/automated-electrophoresis/bioanalyzer-systems**.

**www.agilent.com**

© Agilent Technologies Inc. 2001-2022

Printed in Germany, Edition: 11/2022

Part No: G2938-90037 Rev. E.00

Document No: SD-UF0000031 Rev. E.00
