## Supplementary File 2 for "Transcriptome-wide mapping of small-molecule RNA-binding sites in live cells"

### STAR manual 2.7.11b

Alexander Dobin


January 25, 2024

**7 Output in transcript coordinates. 18**

**8 Counting number of reads per gene. 18**

**9 2-pass mapping. 19**

**10 Merging and mapping of overlapping paired-end reads. 20**

**11 Detection of personal variants overlapping alignments. 20**

**12 WASP filtering of allele specific alignments. 20**

**13 STARconsensus 21**

**14 STARdiploid 21**

**15 Detection of multimapping chimeras. 22**

**16 STARsolo: mapping, demultiplexing and gene quantification for single cell RNA-seq 22**

### 1 Getting started.

#### 1.1 Installation.

STAR source code and binaries can be downloaded from GitHub: named releases from https://github.com/alexdobin/STAR/releases, or the master branch from https://github.com/alexdobin/STAR. The pre-compiled STAR executables are located `bin/` subdirectory. The `static` executables are the easisest to use, as they are statically compiled and are not dependents on external libraries.

To compile STAR from sources run `make` in the source directory for a Linux-like environment, or run `make STARforMac` for Mac OS X. This will produce the executable `'STAR'` inside the source directory.

##### 1.1.1 Installation - in depth and troubleshooting.

STAR is compiled with gcc c++ compiler and depends only on standard gcc libraries. Some generic instructions on installing correct gcc environments are given below.

**Ubuntu.**

```
$ sudo apt-get update
$ sudo apt-get install g++
$ sudo apt-get install make
```

**Red Hat, CentOS, Fedora.**

```
$ sudo yum update
$ sudo yum install make
$ sudo yum install gcc-c++
$ sudo yum install glibc-static
```

**SUSE.**

```
$ sudo zypper update
$ sudo zypper in gcc gcc-c++
```

**Mac OS X.**

Current versions of Mac OS X Xcode are shipped with Clang replacing the standard gcc compiler. Presently, standard Clang does not support OpenMP which creates problems for STAR compilation. One option to avoid this problem is to install gcc (preferably using `homebrew` package manager). Another option is to add OpenMP functionality to Clang.

#### 1.2 Basic workflow.

Basic STAR workflow consists of 2 steps:

1. Generating genome indexes files (see Section 2. Generating genome indexes.
   In this step user supplied the reference genome sequences (FASTA files) and annotations (GTF file), from which STAR generate genome indexes that are utilized in the

2nd (mapping) step. The genome indexes are saved to disk and need only be generated **once** for each genome/annotation combination. A limited collection of STAR genomes is available from http://labshare.cshl.edu/shares/gingeraslab/www-data/dobin/STAR/STARgenomes/, however, it is strongly recommended that users generate their own genome indexes with most up-to-date assemblies and annotations.

2. Mapping reads to the genome (see Section 3. Running mapping jobs).
In this step user supplies the genome files generated in the 1st step, as well as the RNA-seq reads (sequences) in the form of FASTA or FASTQ files. STAR maps the reads to the genome, and writes several output files, such as alignments (SAM/BAM), mapping summary statistics, splice junctions, unmapped reads, signal (wiggle) tracks etc. Output files are described in Section 5. Output files. Mapping is controlled by a variety of input parameters (options) that are described in brief in Section 3. Running mapping jobs, and in more detail in Section 17. Description of all options.

STAR command line has the following format:

```
STAR --option1-name option1-value(s)--option2-name option2-value(s) ...
```

If an option can accept multiple values, they are separated by spaces, and in a few cases - by commas.

### 2 Generating genome indexes.

#### 2.1 Basic options.

The basic options to generate genome indices are as follows:

```
--runThreadN NumberOfThreads
--runMode genomeGenerate
--genomeDir /path/to/genomeDir
--genomeFastaFiles /path/to/genome/fasta1 /path/to/genome/fasta2 ...
--sjdbGTFfile /path/to/annotations.gtf
--sjdbOverhang ReadLength-1
```

`--runThreadN` option defines the number of threads to be used for genome generation, it has to be set to the number of available cores on the server node.

`--runMode genomeGenerate` option directs STAR to run genome indices generation job.

`--genomeDir` specifies path to the directory (henceforth called "genome directory" where the genome indices are stored. This directory has to be created (with `mkdir`) before STAR run and needs to have writing permissions. The file system needs to have at least 100GB of disk space available for a typical mammalian genome. It is recommended to remove all files from the genome directory before running the genome generation step. This directory path will have to be supplied at the mapping step to identify the reference genome.

`--genomeFastaFiles` specifies one or more FASTA files with the genome reference sequences. Multiple reference sequences (henceforth called "chromosomes") are allowed for each fasta file. You can rename the chromosomes' names in the chrName.txt keeping the order of the chromosomes in the file: the names from this file will be used in all output alignment files (such as .sam). The tabs are not allowed in chromosomes' names, and spaces are not recommended.

`--sjdbGTFfile` specifies the path to the file with annotated transcripts in the standard GTF format. STAR will extract splice junctions from this file and use them to greatly improve accuracy of the mapping. While this is optional, and STAR can be run without annotations, using annotations is **highly recommended** whenever they are available. Starting from 2.4.1a, the annotations can also be included on the fly at the mapping step.

`--sjdbOverhang` specifies the length of the genomic sequence around the annotated junction to be used in constructing the splice junctions database. Ideally, this length should be equal to the *ReadLength-1*, where *ReadLength* is the length of the reads. For instance, for Illumina 2x100b paired-end reads, the ideal value is 100-1=99. In case of reads of varying length, the ideal value is *max(ReadLength)-1*. **In most cases, the default value of 100 will work as well as the ideal value.**

Genome files comprise binary genome sequence, suffix arrays, text chromosome names/lengths, splice junctions coordinates, and transcripts/genes information. Most of these files use internal STAR format and are not intended to be utilized by the end user. It is strongly **not recommended** to change any of these files with one exception: you can rename the chromosome names in the chrName.txt while keeping the order of the chromosomes in this file: the chromosome names from this file will be used in all output files (e.g. SAM/BAM).

#### 2.2 Advanced options.

##### 2.2.1 Which chromosomes/scaffolds/patches to include?

It is strongly recommended to include major chromosomes (e.g., for human chr1-22,chrX,chrY,chrM,) as well as un-placed and un-localized scaffolds. Typically, un-placed/un-localized scaffolds add just a few MegaBases to the genome length, however, a substantial number of reads may map to ribosomal RNA (rRNA) repeats on these scaffolds. These reads would be reported as unmapped if the scaffolds are not included in the genome, or, even worse, may be aligned to wrong loci on the chromosomes. Generally, patches and alternative haplotypes should **not** be included in the genome.

Examples of acceptable genome sequence files:

- **ENSEMBL:** files marked with .dna.primary.assembly, such as: ftp://ftp.ensembl.org/pub/release-77/fasta/homo_sapiens/dna/Homo_sapiens.GRCh38.dna.primary_assembly.fa.gz
- **GENCODE:** files marked with PRI (primary). Strongly recommended for mouse and human: http://www.gencodegenes.org/.

##### 2.2.2 Which annotations to use?

The use of the most comprehensive annotations for a given species is strongly recommended. Very importantly, chromosome names in the annotations GTF file have to match chromosome names in the FASTA genome sequence files. For example, one can use ENSEMBL FASTA files with ENSEMBL GTF files, and UCSC FASTA files with UCSC FASTA files. However, since UCSC uses `chr1, chr2, ...` naming convention, and ENSEMBL uses `1, 2, ...` naming, the ENSEMBL and UCSC FASTA and GTF files cannot be mixed together, unless chromosomes are renamed to match between the FASTA anf GTF files.

###### 2.2.3 Annotations in GFF format.

In addition to the aforementioned options, for GFF3 formatted annotations you need to use `--sjdbGTFtagExonParentTranscript Parent`. In general, for `--sjdbGTFfile` files STAR only processes lines which have `--sjdbGTFfeatureExon` (=`exon` by default) in the 3rd field (column). The exons are assigned to the transcripts using parent-child relationship defined by the `--sjdbGTFtagExonParentTranscript` (=`transcript_id` by default) GTF/GFF attribute.

###### 2.2.4 Using a list of annotated junctions.

STAR can also utilize annotations formatted as a list of splice junctions coordinates in a text file: `--sjdbFileChrStartEnd` */path/to/sjdbFile.txt*. This file should contains 4 columns separated by tabs:

`Chr \tab Start \tab End \tab Strand=+/-/.`

Here Start and End are first and last bases of the introns (1-based chromosome coordinates). This file can be used in addition to the `--sjdbGTFfile`, in which case STAR will extract junctions from both files.

Note, that the `--sjdbFileChrStartEnd` file can contain duplicate (identical) junctions, STAR will collapse (remove) duplicate junctions.

###### 2.2.5 Very small genome.

For small genomes, the parameter `--genomeSAindexNbases` **must** to be scaled down, with a typical value of `min(14, log2(GenomeLength)/2 - 1)`. For example, for 1 megaBase genome, this is equal to 9, for 100 kiloBase genome, this is equal to 7.

###### 2.2.6 Genome with a large number of references.

If you are using a genome with a large (>5,000) number of references (chromosomes/scaffolds), you may need to reduce the `--genomeChrBinNbits` to reduce RAM consumption. The following scaling is recommended: `--genomeChrBinNbits` = `min(18,log2[max(GenomeLength/NumberOfReferences,ReadLength)])`. For example, for 3 gigaBase genome with 100,000 chromosomes/scaffolds, this is equal to 15.

### 3 Running mapping jobs.

#### 3.1 Basic options.

The basic options to run a mapping job are as follows:

```
--runThreadN NumberOfThreads
--genomeDir /path/to/genomeDir
--readFilesIn /path/to/read1 [/path/to/read2]
```

`--genomeDir` specifies path to the genome directory where genome indices where generated (see Section 2. Generating genome indexes).

`--readFilesIn` name(s) (with path) of the files containing the sequences to be mapped (e.g. RNA-seq FASTQ files). If using Illumina paired-end reads, the ***read1*** and ***read2*** files have to be supplied. STAR can process both FASTA and FASTQ files. Multi-line (i.e. sequence split in multiple lines) FASTA (but not FASTQ) files are supported.

If the read files are compressed, use the `--readFilesCommand` ***UncompressionCommand*** option, where ***UncompressionCommand*** is the un-compression command that takes the file name as input parameter, and sends the uncompressed output to stdout. For example, for gzipped files (*.gz) use `--readFilesCommand zcat` OR `--readFilesCommand gunzip -c`. For bzip2-compressed files, use `--readFilesCommand bunzip2 -c`.

#### 3.2 Mapping multiple files in one run.

Multiple samples can be mapped in one run with a single output. This is equivalent to concatenating the read files before mapping, except that distinct read groups can be used in `--outSAMattrRGline` command to keep track of reads from different files. For single-end reads use a comma separated list (no spaces around commas), e.g.:

`--readFilesIn sample1.fq,sample2.fq,sample3.fq`

For paired-end reads, use comma separated list for read1, followed by space, followed by comma separated list for read2, e.g.:

`--readFilesIn s1read1.fq,s2read1.fq,s3read1.fq s1read2.fq,s2read2.fq,s3read2.fq`

For multiple read files, the corresponding read groups can be supplied with space/comma/space-separated list in `--outSAMattrRGline`, e.g.

`--outSAMattrRGline ID:sample1 , ID:sample2 , ID:sample3`

Note that this list is separated by commas surrounded by spaces (unlike `--readFilesIn` list).

Another option for mapping multiple reads files, especially convenient for a very large number of files, is to create a file manifest and supply it in `--readFilesManifest /path/to/manifest.tsv`. The manifest file should contain 3 tab-separated columns. For paired-end reads:

`read1-file-name` *tab* `read2-file-name` *tab* `read-group-line`

For single-end reads, the 2nd column should contain the dash -:

`read1-file-name` *tab* `-` *tab* `read-group-line`

Spaces, but not tabs are allowed in the file names. If read-group-line does not start with ID:, it can only contain one ID field, and ID: will be added to it. If read-group-line starts with ID:, it can contain several fields separated by *tab*, and all the fields will be copied verbatim into SAM @RG header line.

#### 3.3 Advanced options.

There are many advanced options that control STAR mapping behavior. All options are briefly described in the Section Section 17. Description of all options.

##### 3.3.1 Using annotations at the mapping stage.

Since 2.4.1a, the annotations can be included on the fly at the mapping step, without including them at the genome generation step. You can specify `--sjdbGTFfile` ***/path/to/ann.gtf*** and/or `--sjdbFileChrStartEnd` ***/path/to/sj.tab***, as well as `--sjdbOverhang`, and any other `--sjdb*` options. The genome indices can be generated with or without another set of annotations/junctions. In the latter case the new junctions will added to the old ones. STAR will insert the junctions into genome indices on the fly before mapping, which takes 1 2 minutes. The on the fly genome indices can be saved (for reuse) with `--sjdbInsertSave All`, into *_STARgenome* directory inside the current run directory.

##### 3.3.2 ENCODE options

An example of ENCODE standard options for long RNA-seq pipeline is given below:

`--outFilterType` BySJout
reduces the number of "spurious" junctions

`--outFilterMultimapNmax` 20
max number of multiple alignments allowed for a read: if exceeded, the read is considered unmapped

`--alignSJoverhangMin` 8
minimum overhang for unannotated junctions

`--alignSJDBoverhangMin` 1
minimum overhang for annotated junctions

`--outFilterMismatchNmax` 999
maximum number of mismatches per pair, large number switches off this filter

`--outFilterMismatchNoverReadLmax` 0.04
max number of mismatches per pair relative to read length: for 2x100b, max number of mismatches is 0.04*200=8 for the paired read

`--alignIntronMin` 20
minimum intron length

`--alignIntronMax` 1000000
maximum intron length

`--alignMatesGapMax` 1000000
maximum genomic distance between mates

#### 3.4 Using shared memory for the genome indexes.

The `--genomeLoad` option controls how the genome is loaded into memory. By default, `--genomeLoad` *NoSharedMemory*, shared memory is not used.

With `--genomeLoad` *LoadAndKeep*, STAR loads the genome as a standard Linux shared memory piece. The genomes are identified by their unique directory paths. Before loading the genome, STAR checks if the genome has already been loaded into the shared memory. If the genome has not been loaded, STAR will load it and will keep it in memory even after STAR job finishes. The genome will be shared with all the other STAR jobs. You can remove the genome from the shared memory running STAR with --genomeLoad *Remove*. The shared memory piece will be physically removed only after all STAR jobs attached to it complete. With --genomeLoad *LoadAndRemove*, STAR will load genome in the shared memory, and mark it for removal, so that the genome will be removed from the shared memory once all STAR jobs using it exit. --genomeLoad *LoadAndExit*, STAR will load genome in the shared memory, and immediately exit, keeping the genome loaded in the shared memory for the future runs.

If you need to check or remove shared memory pieces manually, use the standard Linux command ipcs and ipcrm. If the genome residing in shared memory is not used for a long time it may get paged out of RAM which will slow down STAR runs considerably. It is strongly recommended to regularly re-load (i.e. remove and load again) the shared memory genomes.

Many standard Linux distributions do not allow large enough shared memory blocks. You can fix this issue if you have root privileges, or ask you system administrator to do it. To enable the shared memory modify or add the following lines to /etc/sysctl.conf:

```
kernel.shmmax = Nmax
kernel.shmall = Nall
```

$Nmax$, $Nall$ numbers should be chosen as follows:
$Nmax > GenomeIndexSize = Genome + SA + SAindex$ ( 31000000000 for human genome)
$Nall > GenomeIndexSize/PageSize$
where PageSize is typically 4096 (this can be checked with `getconf PAGE_SIZE`). Then run:

```
/sbin/sysctl -p
```

This will increase the allowed shared memory blocks to 31GB, enough for human or mouse genome.

### 4 Output filtering.

#### 4.1 Multimappers.

The output of multimappers (i.e. reads mapping to multiple loci) is controlled by --outFilterMultimapNmax N. By default, N=10. If a read maps to ≤N loci, it will be output, otherwise, it will be considered as unmapped and reported as "Multimapping: mapped to too many loci" in the `Log.final.out` summary statistics file.

The detection of multimappers is controlled by --winAnchorMultimapNmax option, =50 by default. This parameter should be set to at least the number of multimapping loci, i.e. --winAnchorMultimapNmax ≥ --outFilterMultimapNmax. Note that this parameter also controls the overall sensitivity of mapping: increasing it will change (improve) the mapping of unique mappers as well, though at the cost of slower speed.

### 5 Output files.

STAR produces multiple output files. All files have standard name, however, you can change the file prefixes using --outFileNamePrefix */path/to/output/dir/prefix*. By default, this parameter is `./`, i.e. all output files are written in the current directory.

#### 5.1 Log files.

Log.out: main log file with a lot of detailed information about the run. This file is most useful for troubleshooting and debugging.

Log.progress.out: reports job progress statistics, such as the number of processed reads, % of mapped reads etc. It is updated in 1 minute intervals.

Log.final.out: summary mapping statistics after mapping job is complete, very useful for quality control. The statistics are calculated for each read (single- or paired-end) and then summed or averaged over all reads. Note that STAR counts a paired-end read as one read, (unlike the samtools flagstat/idxstats, which count each mate separately). Most of the information is collected about the UNIQUE mappers (unlike samtools flagstat/idxstats which does not separate unique or multi-mappers). Each splicing is counted in the numbers of splices, which would correspond to summing the counts in SJ.out.tab. The mismatch/indel error rates are calculated on a per base basis, i.e. as total number of mismatches/indels in all unique mappers divided by the total number of mapped bases.

#### 5.2 SAM.

Aligned.out.sam - alignments in standard SAM format.

##### 5.2.1 Multimappers.

The number of loci Nmap a read maps to is given by NH:i:Nmap field. Value of 1 corresponds to unique mappers, while values >1 corresponds to multi-mappers. HI attrbiutes enumerates multiple alignments of a read starting with 1 (this can be changed with the --outSAMattrIHstart - setting it to 0 may be required for compatibility with downstream software such as Cufflinks).

The mapping quality MAPQ (column 5) is 255 for uniquely mapping reads, and int(-10*log10(1-1/Nmap)) for multi-mapping reads. This scheme is same as the one used by TopHat and is compatible with Cufflinks. The default MAPQ=255 for the unique mappers maybe changed with --outSAMmapqUnique parameter (integer 0 to 255) to ensure compatibility with downstream tools such as GATK.

For multi-mappers, all alignments except one are marked with 0x100 (secondary alignment) in the FLAG (column 2 of the SAM). The unmarked alignment is selected from the best ones (i.e. highest scoring). This default behavior can be changed with --outSAMprimaryFlag AllBestScore option, that will output all alignments with the best score as primary alignments (i.e. 0x100 bit in the FLAG unset).

By default, the order of the multi-mapping alignments for each read is not truly random. The --outMultimapperOrder Random option outputs multiple alignments for each read in random order, and also also randomizes the choice of the primary alignment from the highest scoring alignments. Parameter --runRNGseed can be used to set the random generator seed. With this option, the ordering of multi-mapping alignments of each read, and the choice of the primary alignment will vary from run to run, unless only one thread is used and the seed is kept constant.

The --outSAMmultNmax parameter limits the number of output alignments (SAM lines) for multimappers. For instance, --outSAMmultNmax 1 will output exactly one SAM line for each mapped read. Note that `NH:i:` tag in STAR will still report the actual number of loci that the reads map to, while the the number of reported alignments for a read in the SAM file is `min(NH,--outSAMmultNMax)`. If `--outSAMmultNmax` is equal to `-1`, all the alignments are output according to the order specified in `--outMultimapperOrder` option. If `--outSAMmultNmax` is not equal to -1, than top-scoring alignments will always be output first, even for the default `--outMultimapperOrder` `Old_2.4` option.

###### 5.2.2 SAM attributes.

The SAM attributes can be specified by the user using `--outSAMattributes` *`A1 A2 A3 ...`* option which accept a list of 2-character SAM attributes. The attributes can be listed in any order, and will be recorded in that order in the SAM file. By default, STAR outputs `NH HI AS nM` attributes.

**Presets:**

`None` : No SAM attributes

`Standard` : NH HI AS nM

`All` : NH HI AS nM NM MD jM jI MC ch

**Alignment:**

`NH` : number of loci the reads maps to: $= 1$ for unique mappers, $> 1$ for multimappers. Standard SAM tag.

`HI` : multiple alignment index, starts with –outSAMattrIHstart ($= 1$ by default). Standard SAM tag.

`AS` : local alignment score, $+1/-1$ for matches/mismateches, score* penalties for indels and gaps. For PE reads, total score for two mates. Standard SAM tag.

`NM` : edit distance to the reference (number of mismatched + inserted + deleted bases) for each mate. Standard SAM tag.

`nM` : number of mismatches per (paired) alignment, not to be confused with `NM`, which is the number of mismatches+indels in each mate.

`jM:B:c,M1,M2,...` : intron motifs for all junctions (i.e. N in CIGAR): 0: non-canonical; 1: GT/AG, 2: CT/AC, 3: GC/AG, 4: CT/GC, 5: AT/AC, 6: GT/AT. If splice junctions database is used, and a junction is annotated, 20 is added to its motif value.

`MD` : string encoding mismatched and deleted reference bases (see standard SAM specifications). Standard SAM tag.

`jI:B:I,Start1,End1,Start2,End2,...` : Start and End of introns for all junctions (1-based).

`jM jI` : attributes require samtools 0.1.18 or later, and were reported to be incompatible with some downstream tools such as Cufflinks.

**Variation:**

`vA` : variant allele.

`vG` : genomic coordinate of the variant overlapped by the read.

`vW` : WASP filtering tag, see detailed description in Section 12. Requires `--waspOutputMode SAMtag`.

**STARsolo:**

`CR CY UR UY` : sequences and quality scores of cell barcodes and UMIs for the solo* demultiplexing, not error corrected.

`GX GN` : gene ID and name.

`CB UB` : error-corrected cell barcodes and UMIs for solo* demultiplexing. Requires `--outSAMtype BAM SortedByCoordinate`.

`sM` : assessment of CB and UMI.

`sS` : sequence of the entire barcode (CB,UMI,adapter...).

`sQ` : quality of the entire barcode.

**Unmapped reads:**

`uT` : for unmapped reads, reason for not mapping:

0 : no acceptable seed/windows, "Unmapped other" in the Log.final.out
1 : best alignment shorter than min allowed mapped length, "Unmapped: too short" in the Log.final.out
2 : best alignment has more mismatches than max allowed number of mismatches, "Unmapped: too many mismatches" in the Log.final.out
3 : read maps to more loci than the max number of multimappng loci, "Multimapping: mapped to too many loci" in the Log.final.out
4 : unmapped mate of a mapped paired-end read

##### 5.2.3 Compatibility with Cufflinks/Cuffdiff.

For unstranded RNA-seq data, Cufflinks/Cuffdiff require spliced alignments with `XS` strand attribute, which STAR will generate with `--outSAMstrandField intronMotif` option. As required, the XS strand attribute will be generated for all alignments that contain splice junctions. The spliced alignments that have undefined strand (i.e. containing only non-canonical unannotated junctions) will be suppressed.

If you have stranded RNA-seq data, you do not need to use any specific STAR options. Instead, you need to run Cufflinks with the library option `--library-type` options. For example, `cufflinks ... --library-type fr-firststrand` should be used for the "standard" dUTP protocol, including Illumina's stranded Tru-Seq. This option has to be used only for Cufflinks runs and not for STAR runs.

In addition, it is recommended to remove the non-canonical junctions for Cufflinks runs using `--outFilterIntronMotifs RemoveNoncanonical`.

#### 5.3 Unsorted and sorted-by-coordinate BAM.

STAR can output alignments directly in binary BAM format, thus saving time on converting SAM files to BAM. It can also sort BAM files by coordinates, which is required by many downstream applications.

`--outSAMtype BAM Unsorted`
output unsorted `Aligned.out.bam` file. The paired ends of an alignment are always adjacent, and multiple alignments of a read are adjacent as well. This "unsorted" file can be directly used with downstream software such as `HTseq`, without the need of name sorting. The order of the reads will match that of the input FASTQ(A) files only if one thread is used `--runThread 1`, and `--outFilterType --BySJout` is **not** used.

`--outSAMtype BAM SortedByCoordinate`
output sorted by coordinate `Aligned.sortedByCoord.out.bam` file, similar to `samtools sort` command. If this option causes problems, it is recommended to reduce `--outBAMsortingThreadN` from the default 6 to lower values (as low as 1).

`--outSAMtype BAM Unsorted SortedByCoordinate`
output both unsorted and sorted files.

#### 5.4 Unmapped reads.

Unmapped reads can be output into the SAM/BAM `Aligned.*` file(s) with `--outSAMunmapped Within` option. `--outSAMunmapped Within KeepPairs` will (redundantly) record unmapped mate for each alignment, and, in case of unsorted output, keep it adjacent to its mapped mate (this only affects multi-mapping reads). `uT` SAM tag indicates reason for not mapping:

0 : no acceptable seed/windows, "Unmapped other" in the Log.final.out
1 : best alignment shorter than min allowed mapped length, "Unmapped: too short" in the Log.final.out
2 : best alignment has more mismatches than max allowed number of mismatches, "Unmapped: too many mismatches" in the Log.final.out
3 : read maps to more loci than the max number of multimappng loci, "Multimapping: mapped to too many loci" in the Log.final.out
4 : unmapped mate of a mapped paired-end read

`--outReadsUnmapped Fastx` will output unmapped reads into separate file(s) `Unmapped.out.mate1[2]`, formatted the same way as input read files (i.e. FASTQ or FASTA). For paired-end reads, if a read maps as a whole, but one of the mates does not map, both mates will also be output in `Unmapped.out.mate1/2` files. To indicate the mapping status of the read mates, the following tags are appended to the read name:
00: mates were not mapped;
10: 1st mate mapped, 2nd unmapped
01: 1st unmapped, 2nd mapped

#### 5.5 Splice junctions.

`SJ.out.tab` contains high confidence collapsed splice junctions in tab-delimited format. Note that STAR defines the junction start/end as intronic bases, while many other software define them as exonic bases. The columns have the following meaning:

column 1: chromosome

column 2: first base of the intron (1-based)

column 3: last base of the intron (1-based)

column 4: strand (0: undefined, 1: +, 2: -)

column 5: intron motif: 0: non-canonical; 1: GT/AG, 2: CT/AC, 3: GC/AG, 4: CT/GC, 5: AT/AC, 6: GT/AT

column 6: 0: unannotated, 1: annotated in the splice junctions database. Note that in 2-pass mode, junctions detected in the 1st pass are reported as annotated, in addition to annotated junctions from GTF.

column 7: number of uniquely mapping reads crossing the junction

column 8: number of multi-mapping reads crossing the junction

column 9: maximum spliced alignment overhang

The filtering for this output file is controlled by the `--outSJfilter*` parameters, as described in Section 17.18. Output Filtering: Splice Junctions.

### 6 Chimeric and circular alignments.

To switch on detection of chimeric (fusion) alignments (in addition to normal mapping), `--chimSegmentMin` should be set to a positive value. Each chimeric alignment consists of two "segments". Each segment is non-chimeric on its own, but the segments are chimeric to each other (i.e. the segments belong to different chromosomes, or different strands, or are far from each other). Both segments may contain splice junctions, and one of the segments may contain portions of both mates. `--chimSegmentMin` parameter controls the minimum mapped length of the two segments that is allowed. For example, if you have 2x75 reads and used `--chimSegmentMin` 20, a chimeric alignment with 130b on one chromosome and 20b on the other will be output, while 135 + 15 won't be.

#### 6.1 STAR-Fusion.

STAR-Fusion is a software package for detecting fusion transcript from STAR chimeric output. It is developed and maintained by Brian Haas (@Broad Institute), whose effort was inspired by earlier work done by Nicolas Stransky in the landmark publication "The landscape of kinase fusions in cancer" by Stransky et al., Nat Commun 2014, in addition to very nice work done by Daniel Nicorici with his FusionCatcher software. Please visit its GitHub page for instructions and documentation: https://github.com/STAR-Fusion/STAR-Fusion.

#### 6.2 Chimeric alignments in the main BAM files.

Chimeric alignments can be included together with normal alignments in the main (sorted or unsorted) BAM file(s) using `--chimOutType WithinBAM`. In these files, formatting of chimeric alignments follows the latest SAM/BAM specifications.

#### 6.3 Chimeric alignments in `Chimeric.out.sam` .

With `--chimOutType SeparateSAMold` STAR will output normal alignments into `Aligned.*.sam/bam`, and will output chimeric alignments into a separate file `Chimeric.out.sam`. Note that this option will be deprecated in the future, and the `--chimOutType WithinBAM` is strongly recommended. Some reads may be output to both normal SAM/BAM files, and `Chimeric.out.sam` for the following reason. STAR will output a non-chimeric alignment into `Aligned.out.sam` with soft-clipping a portion of the read. If this portion is long enough, and it maps well and uniquely somewhere else in the genome, there will also be a chimeric alignment output into `Chimeric.out.sam`. For instance, if you have a paired-end read where the second mate can be split chimerically into 70 and 30 bases. The 100b of the first mate + 70b of the 2nd mate map non-chimerically,and the mapping length/score are big enough, so they will be output into `Aligned.out.sam` file. At the same time, the chimeric segments 100-mate1 + 70-mate2 and 30-mate2 will be output into `Chimeric.out.sam`.

#### 6.4 Chimeric alignments in `Chimeric.out.junction`

By default, or with `--chimOutType Junctions`, STAR will generate `Chimeric.out.junction` file which maybe more convenient for downstream analysis. The format of this file is as follows. Every line contains one chimerically aligned read, e.g.:

```
chr22   23632601        +       chr9    133729450       +       1       0       0
SINATRA-0006:3:3:6387:5665#0    23632554   47M29S  133729451       47S29M40p76M
```

The first 9 columns give information about the chimeric junction:

column 1: **chr_donorA** : chromosome of the donor

column 2: **brkpt_donorA** : first base of the intron of the donor (1-based)

column 3: **strand_donorA** : strand of the donor

column 4: **chr_acceptorB** : chromosome of the acceptor

column 5: **brkpt_acceptorB** : first base of the intron of the acceptor (1-based)

column 6: **strand_acceptorB** : strand of the acceptor

column 7: **junction_type** : -1=encompassing junction (between the mates), 1=GT/AG, 2=CT/AC

column 8: **repeat_left_lenA** : repeat length to the left of the junction

column 9: **repeat_right_lenB** : repeat length to the right of the junction

Columns 10-14 describe the alignments of the two chimeric segments, it is SAM like. Alignments are given with respect to the (+) strand

column 10: **read_name** : name of the RNA-seq fragment

column 11: **start_alnA** : first base of the first segment (on the + strand)

column 12: **cigar_alnA** : CIGAR of the first segment

column 13: **start_alnB** : first base of the second segment

column 14: **cigar_alnB** : CIGAR of the second segment

Columns 15-20 provide alignment score information and relevant metadata. These columns are only output for multimapping chimeriuc algorithm `--chimMultimapNmax` `>0`.

column 15: **num_chim_aln** : number of sufficiently scoring chimeric alignments reported for this RNA-seq fragment.

column 16: **max_poss_aln_score** : maximum possible alignment score for this fragment's read(s).

column 17: **non_chim_aln_score** : best non-chimeric alignment score

column 18: **this_chim_aln_score** : score for this individual chimeric alignment

column 19: **bestall_chim_aln_score** : the highest chimeric alignment score encountered for this RNA-seq fragment among the **num_chim_aln** reported chimeric alignments.

column 20: **PEmerged_bool** : boolean indicating that overlapping PE reads were first merged into a single contiguous sequence before alignment.

column 21: **readgrp** : read group assignment for the read as indicated in the BAM file

Unlike standard SAM, both mates are recorded in one line here. The gap of length `L` between the mates is marked by the `p` in the CIGAR string. If the mates overlap, `L<0`.

For strand definitions, when aligning paired end reads, the sequence of the second mate is reverse complemented.

For encompassing junctions, i.e. junction type: -1=junction is between the mates, columns 2 and 5 represent the bounds on the chimeric junction loci. For the 1st mate, it will be the genomic base following the last 3' mapped base. For the 2nd mate (which is reverse complemented to have the same orientation as 1st mate), it will be the genomic base preceding the 5' mapped base. For example, if there is a chimeric junction that connects chr1/+strand/base1000 to chr2/+strand/base2000, and read 1 maps to chr1/+strand/bases800-900, and read 2 (after reverse complementing) maps to chr2/+strand/bases2100-2200, then columns 2 and 5 will have 901 and 2099.

To filter chimeric junctions and find the number of reads supporting each junction you could use, for example:

```
cat Chimeric.out.junction |
awk '$1!="chrM" && $4!="chrM" && $7>0 && $8+$9<=5 {print $1,$2,$3,$4,$5,$6,$7,$8,$9}' |
sort | uniq -c | sort -k1,1rn
```

This will keep only the canonical junctions with the repeat length less than 5 and will remove chimeras with mitochondrion genome.

When I do it for one of our K562 runs, I get:

| | | | | | | | | | |
|---|---|---|---|---|---|---|---|---|---|
| 181 | chr1 | 144676873 | - | chr1 | 147917466 | + | 1 | 0 | 1 |
| 29 | chr5 | 69515744 | - | chr5 | 34182973 | - | 1 | 3 | 1 |
| 28 | chr1 | 143910077 | - | chr1 | 149459550 | - | 1 | 1 | 0 |
| 27 | chr22 | 23632601 | + | chr9 | 133729450 | + | 1 | 0 | 0 |
| 20 | chr12 | 90313405 | - | chr21 | 40684813 | - | 1 | 2 | 0 |
| 20 | chr22 | 23632601 | + | chr9 | 133655755 | + | 1 | 0 | 1 |
| 20 | chr9 | 123636256 | - | chr9 | 123578959 | + | 1 | 1 | 4 |
| 15 | chr16 | 85589970 | + | chr6 | 16762582 | + | 1 | 3 | 2 |
| 15 | chr3 | 197348574 | - | chr3 | 195392936 | + | 1 | 1 | 0 |
| 14 | chr18 | 39584506 | + | chr18 | 39560613 | - | 1 | 2 | 0 |

Note that line 4 and 6 here are BCR/ABL fusions. You would need to filter these junctions further to see which of them connect known but not homologous genes.

### 7 Output in transcript coordinates.

With `--quantMode` `TranscriptomeSAM` option STAR will output alignments translated into transcript coordinates in the `Aligned.toTranscriptome.out.bam` file (in addition to alignments in genomic coordinates in `Aligned.*.sam/bam` files). These transcriptomic alignments can be used with various transcript quantification software that require reads to be mapped to transcriptome, such as RSEM or eXpress. For example, RSEM command line would look as follows:

```
rsem-calculate-expression ...  --bam Aligned.toTranscriptome.out.bam
/path/to/RSEM/reference RSEM
```

Note, that STAR first aligns reads to entire genome, and only then searches for concordance between alignments and transcripts.This approach offers certain advantages compared to the alignment to transcriptome only, by not forcing the alignments to annotated transcripts. Note that `--outFilterMultimapNmax` filter only applies to genomic alignments. If an alignment passes this filter, it is converted to all possible transcriptomic alignments and all of them are output.

By default, the output satisfies RSEM requirements: soft-clipping or indels are not allowed. Use `--quantTranscriptomeBan` `Singleend` to allow insertions, deletions ans soft-clips in the transcriptomic alignments, which can be used by some expression quantification software (e.g. eXpress).

### 8 Counting number of reads per gene.

With `--quantMode` `GeneCounts` option STAR will count number reads per gene while mapping. A read is counted if it overlaps (1nt or more) one and only one gene. Both ends of the paired-end read are checked for overlaps. The counts coincide with those produced by htseq-count with default parameters. This option requires annotations in GTF format (i.e. *gene_id* tag for each exon) specified in `--sjdbGTFfile` at the genome generation step or at the mapping step provided in option. STAR outputs read counts per gene into ReadsPerGene.out.tab file with 4 columns which correspond to different strandedness options:

column 1: gene ID

column 2: counts for unstranded RNA-seq

column 3: counts for the 1st read strand aligned with RNA (htseq-count option -s yes)

column 4: counts for the 2nd read strand aligned with RNA (htseq-count option -s reverse)

Select the output according to the strandedness of your data. Note, that if you have stranded data and choose one of the columns 3 or 4, the other column (4 or 3) will give you the count of antisense reads. With `--quantMode` `TranscriptomeSAM GeneCounts`, and get both the `Aligned.toTranscriptome.out.bam` and `ReadsPerGene.out.tab` outputs.

### 9 2-pass mapping.

For the most sensitive novel junction discovery, it is recommended to run STAR in the 2-pass mode. It does not significantly increase the number of detected novel junctions, but allows to detect more splices reads mapping to novel junctions. The basic idea is to run 1st pass of STAR mapping with the usual parameters, then collect the junctions detected in the first pass, and use them as "annotated" junctions for the 2nd pass mapping.

#### 9.1 Multi-sample 2-pass mapping.

For a study with multiple samples, it is recommended to collect 1st pass junctions from all samples.

1. Run 1st mapping pass for all samples with "usual" parameters. Using annotations is recommended either a the genome generation step, or mapping step.

2. Run 2nd mapping pass for all samples , listing SJ.out.tab files from all samples in `--sjdbFileChrStartEnd` */path/to/sj1.tab /path/to/sj2.tab ....*

#### 9.2 Per-sample 2-pass mapping.

Annotated junctions will be included in both the 1st and 2nd passes. To run STAR 2-pass mapping for each sample separately, use `--twopassMode` `Basic` option. STAR will perform the 1st pass mapping, then it will automatically extract junctions, insert them into the genome index, and, finally, re-map all reads in the 2nd mapping pass. This option can be used with annotations, which can be included either at the run-time (see #1), or at the genome generation step.

`--twopass1readsN` defines the number of reads to be mapped in the 1st pass. The default and most sensitive approach is to set it to -1 (or make it bigger than the number of reads in the sample) - in which case all reads in the input read file(s) are used in the 1st pass. While it can reduce mapping time by $\sim$ 40%, it is not recommended to use a small portion of the reads in the 1st step, since it will significantly reduce sensitivity for the low expressed novel junctions. The idea to use a portion of the reads in the 1st pass was inspired by Kim, Langmead and Salzberg in Nature Methods 12, 357–360 (2015).

#### 9.3 2-pass mapping with re-generated genome.

This is the original 2-pass method which involves genome re-generation step in-between 1st and 2nd passes. Since 2.4.1a, it is recommended to use the on the fly 2-pass options as described above.

1. Run 1st pass STAR for all samples with "usual" parameters. Genome indices generated with annotations are recommended.

2. Collect all junctions detected in the 1st pass by merging `SJ.out.tab` files from all runs. Filter the junctions by removing likelie false positives, e.g. junctions in the mitochondrion genome, or non-canonical junctions supported by a few reads. If you are using annotations, only novel junctions need to be considered here, since annotated junctions will be re-used in the 2nd pass anyway.

3. Use the filtered list of junctions from the 1st pass with `--sjdbFileChrStartEnd` option, together with annotations (via `--sjdbGTFfile` option) to generate the new genome indices for the 2nd pass mapping. This needs to be done only once for all samples.

4. Run the 2nd pass mapping for all samples with the new genome index.

### 10 Merging and mapping of overlapping paired-end reads.

This feature improves mapping accuracy for paired-end libraries with short insert sizes, where many reads have overlapping mates. Importantly, it allows detection of chimeric junction in the overlap region.

STAR will search for an overlap between mates larger or equal to `--peOverlapNbasesMin` bases with proportion of mismatches in the overlap area not exceeding `--peOverlapMMp`. If the overlap is found, STAR will map merge the mates and attempt to map the resulting (single-end) sequence. If requested, the chimeric detection will be performed on the merged-mate sequence, thus allowing chimeric detection in the overlap region. If the score of this alignment higher than the original one, or if a chimeric alignment is found, STAR will report the merged-mate aligment instead of the original one. In the output, the merged-mate aligment will be converted back to paired-end format.

The developmment of this algorithm was supported by Illumina, Inc. Many thanks to June Snedecor, Xiao Chen, and Felix Schlesinger for their extensive help in developing this feature.

### 11 Detection of personal variants overlapping alignments.

Option `--varVCFfile` */path/to/vcf/file* is used to input VCF file with personal variants. Only single nucleotide variants (SNVs) are supported at the moment. Each variant is expected to have a genotype with two alleles. To output variants that overlap alignments, vG and vA have to be added to `--outSAMattributes` list. SAM attribute vG outputs the genomic coordinate of the variant, allowing for identification of the variant. SAM attribute vA outputs which allele is detected in the read: 1 or 2 match one of the genotype alleles, 3 - no match to genotype.

### 12 WASP filtering of allele specific alignments.

This is re-implementation of the original WASP algorithm by Bryce van de Geijn, Graham McVicker, Yoav Gilad and Jonathan K Pritchard. Please cite the original WASP paper: Nature Methods 12, 1061–1063 (2015) https://www.nature.com/articles/nmeth.3582. WASP filtering is activated with `--waspOutputMode` `SAMtag`, which will add `vW` tag to the SAM output: `vW:i:1` means alignment passed WASP filtering, and all other values mean it did not pass:

`vW:i:2` - multi-mapping read
`vW:i:3` - variant base in the read is N (non-ACGT)
`vW:i:4` - remapped read did not map
`vW:i:5` - remapped read multi-maps
`vW:i:6` - remapped read maps to a different locus
`vW:i:7` - read overlaps too many variants

### 13 STARconsensus

STARconsensus allows for mapping RNA-seq reads to consensus genome. It was introduced in STAR 2.7.7a (2020/12/28).

- Provide the VCF file with consensus SNVs and InDels at the genome generation stage with `--genomeTransformVCF Variants.vcf` `--genomeTransformType` `Haploid`. The alternative alleles in this VCF will be inserted to the reference genome to create a "transformed" genome. Both the genome sequence and transcript/gene annotations are transformed.

- At the mapping stage, the reads will be mapped to the tranformed (consensus) genome. The quantification in the transformed annotations can be performed with standard `--quantMode` `TranscriptomeSAM` and/or `GeneCounts` options. If desired, alignments (SAM/BAM) and spliced junctions (SJ.out.tab) can be transformed back to the original (reference) coordinates with `--genomeTransformOutput` `SAM` and/or `SJ`. This is useful if downstream processing relies on reference coordinates.

### 14 STARdiploid

STARdiploid allows for mapping RNA-seq reads to diploid genome. It was introduced in STAR 2.7.11a.

- Provide the VCF file with diploid SNVs and InDels at the genome generation stage with `--genomeTransformVCF Variants.vcf` `--genomeTransformType` `Diploid`. The homozygous and heterozygous alleles from this VCF will be inserted into the reference genome to create a "transformed" genome with two haplotypes. Both the genome sequence and transcript/gene annotations are transformed.

- At the mapping stage, the reads will be mapped to the diploid genome. If desired, alignments (SAM/BAM), spliced junctions (SJ.out.tab) and quantifications can be transformed back to the original (reference) coordinates with `--genomeTransformOutput` `SAM`, `SJ` and/or`Quant` option(s). This is useful if downstream processing relies on reference coordinates. SAM attribute `ha` can be added to the `--outSAMattribute` list to mark which haplotype the alignment belongs to: 1 - haplotype 1, 2 - haplotype 2, 0 - equally good alignment to both haplotypes.

### 15 Detection of multimapping chimeras.

Previous STAR chimeric detection algorithm only detected uniquely mapping chimeras, which reduced its sensitivity in some cases. The new algorithm can detect and output multimapping chimeras. Presently, the only output into Chimeric.out.junction is supported. This algorithm is activated with $> 0$ value in `chimMultimapNmax`, which defines the maximum number of chimeric multi-alignments. The `chimMultimapScoreRange` ($= 1$ by default) parameter defines the score range for multi-mapping chimeras below the best chimeric score, similar to the `outFilterMultimapScoreRange` parameter for normal alignments. The `chimNonchimScoreDropMin` ($= 20$ by default) defines the threshold triggering chimeric detection: the drop in the best non-chimeric alignment score with respect to the read length has to be greater than this value.

### 16 STARsolo: mapping, demultiplexing and gene quantification for single cell RNA-seq

STARsolo is a turnkey solution for analyzing droplet single cell RNA sequencing data (e.g. 10X Genomics Chromium System) built directly into STAR code. STARsolo inputs the raw FASTQ reads files, and performs the following operations:

- error correction and demultiplexing of cell barcodes using user-input whitelist
- mapping the reads to the reference genome using the standard STAR spliced read alignment algorithm
- error correction and collapsing (deduplication) of Unique Molecular Identifiers (UMIa)
- quantification of per-cell gene expression by counting the number of reads per gene

STARsolo output is designed to be a drop-in replacement for 10X CellRanger gene quantification output. It follows CellRanger logic for cell barcode whitelisting and UMI deduplication, and produces nearly identical gene counts in the same format. At the same time STARsolo is 10 times faster than the CellRanger.

STARsolo solo* options can be found in the Section 17.27. For more detailed description, please see https://github.com/alexdobin/STAR/blob/master/docs/STARsolo.md.

#### 16.1 Feature statistics summaries.

Feature statistics summaries are recorded in the *Solo.out/* directory in files *<Feature>.stats* where features are those used in the `--soloFeatures` option, e.g. *Gene.stats*. The following metrics are recorded:

`nNinBarcode`: number of reads with more than 2 Ns in cell barcode (CB)
`nUMIhomopolymer`: number of reads with homopolymer in CB
`nTooMany`: not used at the moment
`nNoMatch`: number of reads with CBs that do not match whitelist even with one mismatch

All of the above reads are discarded from Solo output. Remaining reads are checked for overlap with features (e.g. genes):

**nUnmapped:** number of reads unmapped to the genome

**nNoFeature:** number of reads that map to the genome but do not belong to a feature

**nAmbigFeature:** number of reads that belong to more than one feature

**nAmbigFeatureMultimap:** number of reads that belong to more than one feature and are also multimapping to the genome (this is a subset of the nAmbigFeature)

**nTooMany:** number of reads with ambiguous CB (i.e. CB matches whitelist with one mismatch but with posterior probability ¡0.95)

**nNoExactMatch:** number of reads with CB that matches a whitelist barcode with 1 mismatch, but this whitelist barcode does not get any other reads with exact matches of CB

All of the reads above are output in feature (e.g. gene) / cell count matrices.

**nExactMatch:** number of reads with CB that match the whitelist exactly

**nMatch:** total number of reads that match CB with 0 or 1 mismatches (this is superset of nExactMatch)

**nCellBarcodes:** number of distinct CBs detected

**nUMIs:** number of distinct UMIs detected

These metrics can be grouped into more broad categories:

`nNinBarcode+nUMIhomopolymer+nNoMatch+nTooMany+nNoExactMatch` = number of reads with CBs that do not match whitelist.

`nUnmapped+nAmbigFeature` = number of reads without defined feature (gene)

`nMatch` = number of reads that are output as solo counts

The three categoties above summed together should be equal to the total number of reads.

### 17 Description of all options.

For each STAR version, the most up-to-date information about all STAR parameters can be found in the `parametersDefault` file in the STAR source directory. The parameters in the `parametersDefault`, as well as in the descriptions below, are grouped by function:

Special attention has to be paid to parameters that start with `--out*`, as they control the STAR output.

In particular, `--outFilter*` parameters control the filtering of output alignments which[] you might want to tweak to fit your needs.

Output of "chimeric" alignments is controlled by `--chim*` parameters.

Genome generation is controlled by `--genome*` parameters.

Annotations (splice junction database) are controlled by `--sjdb*` options at the genome generation step.

Tweaking `--score*`, `--align*`, `--seed*`, `--win*` parameters, which requires understanding of the STAR alignment algorithm, is recommended only for advanced users.

Below, allowed parameter values are typed in magenta, and default values - in blue.

#### 17.1 Parameter Files

`--parametersFiles`

default: -

string: name of a user-defined parameters file, ”-”: none. Can only be defined on the command line.

#### 17.2 System

`--sysShell`

default: -

string: path to the shell binary, preferably bash, e.g. /bin/bash.

`-`
: the default shell is executed, typically /bin/sh. This was reported to fail on some Ubuntu systems - then you need to specify path to bash.

#### 17.3 Run Parameters

`--runMode`

default: `alignReads`

string: type of the run.

`alignReads`
: map reads

`genomeGenerate`
: generate genome files

`inputAlignmentsFromBAM`
: input alignments from BAM. Presently only works with –outWigType and –bamRemoveDuplicates options.

`liftOver`
: lift-over of GTF files (–sjdbGTFfile) between genome assemblies using chain file(s) from –genomeChainFiles.

`soloCellFiltering </path/to/raw/count/dir/> </path/to/output/prefix>`
: STARsolo cell filtering (”calling”) without remapping, followed by the path to raw count directory and output (filtered) prefix

`--runThreadN`

default: 1

int: number of threads to run STAR

--runDirPerm

default: User_RWX

string: permissions for the directories created at the run-time.

User_RWX
: user-read/write/execute

All_RWX
: all-read/write/execute (same as chmod 777)

--runRNGseed

default: 777

int: random number generator seed.

#### 17.4 Genome Parameters

--genomeDir

default: ./GenomeDir/

string: path to the directory where genome files are stored (for –runMode alignReads) or will be generated (for –runMode generateGenome)

--genomeLoad

default: NoSharedMemory

string: mode of shared memory usage for the genome files. Only used with –runMode alignReads.

LoadAndKeep
: load genome into shared and keep it in memory after run

LoadAndRemove
: load genome into shared but remove it after run

LoadAndExit
: load genome into shared memory and exit, keeping the genome in memory for future runs

Remove
: do not map anything, just remove loaded genome from memory

NoSharedMemory
: do not use shared memory, each job will have its own private copy of the genome

--genomeFastaFiles

default: -

string(s): path(s) to the fasta files with the genome sequences, separated by spaces. These files should be plain text FASTA files, they *cannot* be zipped.

Required for the genome generation (–runMode genomeGenerate). Can also be used in the mapping (–runMode alignReads) to add extra (new) sequences to the genome (e.g. spike-ins).

`--genomeChainFiles`

default: `-`

string: chain files for genomic liftover. Only used with –runMode liftOver .

`--genomeFileSizes`

default: `0`

uint(s)>0: genome files exact sizes in bytes. Typically, this should not be defined by the user.

`--genomeTransformOutput`

default: `None`

string(s): which output to transform back to original genome

`SAM`
: SAM/BAM alignments

`SJ`
: splice junctions (SJ.out.tab)

`Quant`
: quantifications (from –quantMode option)

`None`
: no transformation of the output

`--genomeChrSetMitochondrial`

default: `chrM M MT`

string(s): names of the mitochondrial chromosomes. Presently only used for STARsolo statistics output/

##### 17.5 Genome Indexing Parameters - only used with –runMode genomeGenerate

--genomeChrBinNbits

default: 18

int: =log2(chrBin), where chrBin is the size of the bins for genome storage: each chromosome will occupy an integer number of bins. For a genome with large number of contigs, it is recommended to scale this parameter as min(18, log2[max(GenomeLength/NumberOfReferences,ReadLength)]).

--genomeSAindexNbases

default: 14

int: length (bases) of the SA pre-indexing string. Typically between 10 and 15. Longer strings will use much more memory, but allow faster searches. For small genomes, the parameter –genomeSAindexNbases must be scaled down to min(14, log2(GenomeLength)/2 - 1).

--genomeSAsparseD

default: 1

int>0: suffux array sparsity, i.e. distance between indices: use bigger numbers to decrease needed RAM at the cost of mapping speed reduction

--genomeSuffixLengthMax

default: -1

int: maximum length of the suffixes, has to be longer than read length. -1 = infinite.

--genomeTransformType

default: None

string: type of genome transformation

- None
  no transformation
- Haploid
  replace reference alleles with alternative alleles from VCF file (e.g. consensus allele)
- Diploid
  create two haplotypes for each chromosome listed in VCF file, for genotypes 1—2, assumes perfect phasing (e.g. personal genome)

--genomeTransformVCF

default: -

string: path to VCF file for genome transformation

#### 17.6 Splice Junctions Database

`--sjdbFileChrStartEnd`

default: -

string(s): path to the files with genomic coordinates (chr <tab> start <tab> end <tab> strand) for the splice junction introns. Multiple files can be supplied and will be concatenated.

`--sjdbGTFfile`

default: -

string: path to the GTF file with annotations

`--sjdbGTFchrPrefix`

default: -

string: prefix for chromosome names in a GTF file (e.g. 'chr' for using ENSMEBL annotations with UCSC genomes)

`--sjdbGTFfeatureExon`

default: exon

string: feature type in GTF file to be used as exons for building transcripts

`--sjdbGTFtagExonParentTranscript`

default: transcript_id

string: GTF attribute name for parent transcript ID (default "transcript_id" works for GTF files)

`--sjdbGTFtagExonParentGene`

default: gene_id

string: GTF attribute name for parent gene ID (default "gene_id" works for GTF files)

`--sjdbGTFtagExonParentGeneName`

default: gene_name

string(s): GTF attribute name for parent gene name

`--sjdbGTFtagExonParentGeneType`

default: gene_type gene_biotype

string(s): GTF attribute name for parent gene type

`--sjdbOverhang`

default: `100`

int>0: length of the donor/acceptor sequence on each side of the junctions, ideally = (mate_length - 1)

`--sjdbScore`

default: `2`

int: extra alignment score for alignments that cross database junctions

`--sjdbInsertSave`

default: `Basic`

string: which files to save when sjdb junctions are inserted on the fly at the mapping step

`Basic`
: only small junction / transcript files

`All`
: all files including big Genome, SA and SAindex - this will create a complete genome directory

#### 17.7 Variation parameters

`--varVCFfile`

default: `-`

string: path to the VCF file that contains variation data. The 10th column should contain the genotype information, e.g. 0/1

#### 17.8 Input Files

`--inputBAMfile`

default: `-`

string: path to BAM input file, to be used with –runMode inputAlignmentsFromBAM

#### 17.9 Read Parameters

--readFilesType

default: Fastx

string: format of input read files

Fastx
: FASTA or FASTQ

SAM SE
: SAM or BAM single-end reads; for BAM use –readFilesCommand samtools view

SAM PE
: SAM or BAM paired-end reads; for BAM use –readFilesCommand samtools view

--readFilesSAMattrKeep

default: All

string(s): for –readFilesType SAM SE/PE, which SAM tags to keep in the output BAM, e.g.: –readFilesSAMtagsKeep RG PL

All
: keep all tags

None
: do not keep any tags

--readFilesIn

default: Read1 Read2

string(s): paths to files that contain input read1 (and, if needed, read2)

--readFilesManifest

default: -

string: path to the "manifest" file with the names of read files. The manifest file should contain 3 tab-separated columns:

paired-end reads: read1_file_name *tab* read2_file_name *tab* read_group_line.

single-end reads: read1_file_name *tab* - *tab* read_group_line.

Spaces, but not tabs are allowed in file names.

If read_group_line does not start with ID:, it can only contain one ID field, and ID: will be added to it.

If read_group_line starts with ID:, it can contain several fields separated by *tab*, and all fields will be be copied verbatim into SAM @RG header line.

`--readFilesPrefix`

default: `-`

string: prefix for the read files names, i.e. it will be added in front of the strings in –readFilesIn

`--readFilesCommand`

default: `-`

string(s): command line to execute for each of the input file. This command should generate FASTA or FASTQ text and send it to stdout

For example: zcat - to uncompress .gz files, bzcat - to uncompress .bz2 files, etc.

`--readMapNumber`

default: `-1`

int: number of reads to map from the beginning of the file

-1: map all reads

`--readMatesLengthsIn`

default: `NotEqual`

string: Equal/NotEqual - lengths of names,sequences,qualities for both mates are the same / not the same. NotEqual is safe in all situations.

`--readNameSeparator`

default: `/`

string(s): character(s) separating the part of the read names that will be trimmed in output (read name after space is always trimmed)

`--readQualityScoreBase`

default: `33`

int>=0: number to be subtracted from the ASCII code to get Phred quality score

##### 17.10 Read Clipping

--clipAdapterType

default: Hamming

string: adapter clipping type

Hamming
: adapter clipping based on Hamming distance, with the number of mismatches controlled by –clip5pAdapterMMp

CellRanger4
: 5p and 3p adapter clipping similar to CellRanger4. Utilizes Opal package by Martin Šošić: https://github.com/Martinsos/opal

None
: no adapter clipping, all other clip* parameters are disregarded

--clip3pNbases

default: 0

int(s): number(s) of bases to clip from 3p of each mate. If one value is given, it will be assumed the same for both mates.

--clip3pAdapterSeq

default: -

string(s): adapter sequences to clip from 3p of each mate. If one value is given, it will be assumed the same for both mates.

polyA
: polyA sequence with the length equal to read length

--clip3pAdapterMMp

default: 0.1

double(s): max proportion of mismatches for 3p adapter clipping for each mate. If one value is given, it will be assumed the same for both mates.

--clip3pAfterAdapterNbases

default: 0

int(s): number of bases to clip from 3p of each mate after the adapter clipping. If one value is given, it will be assumed the same for both mates.

--clip5pNbases

default: 0

int(s): number(s) of bases to clip from 5p of each mate. If one value is given, it will be assumed the same for both mates.

##### 17.11 Limits

--limitGenomeGenerateRAM

default: 31000000000

int>0: maximum available RAM (bytes) for genome generation

--limitIObufferSize

default: 30000000 50000000

int(s)>0: max available buffers size (bytes) for input/output, per thread

--limitOutSAMoneReadBytes

default: 100000

int>0: max size of the SAM record (bytes) for one read. Recommended value: >(2*(LengthMate1+LengthMate2+100)*outFilterMultimapNmax

--limitOutSJoneRead

default: 1000

int>0: max number of junctions for one read (including all multi-mappers)

--limitOutSJcollapsed

default: 1000000

int>0: max number of collapsed junctions

--limitBAMsortRAM

default: 0

int>=0: maximum available RAM (bytes) for sorting BAM. If =0, it will be set to the genome index size. 0 value can only be used with –genomeLoad NoSharedMemory option.

--limitSjdbInsertNsj

default: 1000000

int>=0: maximum number of junctions to be inserted to the genome on the fly at the mapping stage, including those from annotations and those detected in the 1st step of the 2-pass run

--limitNreadsSoft

default: -1

int: soft limit on the number of reads

##### 17.12 Output: general

--outFileNamePrefix

default: ./

string: output files name prefix (including full or relative path). Can only be defined on the command line.

--outTmpDir

default: -

string: path to a directory that will be used as temporary by STAR. All contents of this directory will be removed!

-
: the temp directory will default to outFileNamePrefix_STARtmp

--outTmpKeep

default: None

string: whether to keep the temporary files after STAR runs is finished

None
: remove all temporary files

All
: keep all files

--outStd

default: Log

string: which output will be directed to stdout (standard out)

Log
: log messages

SAM
: alignments in SAM format (which normally are output to Aligned.out.sam file), normal standard output will go into Log.std.out

BAM_Unsorted
: alignments in BAM format, unsorted. Requires –outSAMtype BAM Unsorted

BAM_SortedByCoordinate
: alignments in BAM format, sorted by coordinate. Requires –outSAMtype BAM SortedByCoordinate

BAM_Quant
: alignments to transcriptome in BAM format, unsorted. Requires –quantMode TranscriptomeSAM

--outReadsUnmapped

default: None

string: output of unmapped and partially mapped (i.e. mapped only one mate of a paired end read) reads in separate file(s).

None
: no output

Fastx
: output in separate fasta/fastq files, Unmapped.out.mate1/2

--outQSconversionAdd

default: 0

int: add this number to the quality score (e.g. to convert from Illumina to Sanger, use -31)

--outMultimapperOrder

default: Old_2.4

string: order of multimapping alignments in the output files

Old_2.4
: quasi-random order used before 2.5.0

Random
: random order of alignments for each multi-mapper. Read mates (pairs) are always adjacent, all alignment for each read stay together. This option will become default in the future releases.

#### 17.13 Output: SAM and BAM

--outSAMtype

default: SAM

strings: type of SAM/BAM output

1st word:

BAM
: output BAM without sorting

SAM
: output SAM without sorting

None
: no SAM/BAM output

2nd, 3rd:

`Unsorted`
: standard unsorted

`SortedByCoordinate`
: sorted by coordinate. This option will allocate extra memory for sorting which can be specified by –limitBAMsortRAM.

`--outSAMmode`

default: `Full`

string: mode of SAM output

`None`
: no SAM output

`Full`
: full SAM output

`NoQS`
: full SAM but without quality scores

`--outSAMstrandField`

default: `None`

string: Cufflinks-like strand field flag

`None`
: not used

`intronMotif`
: strand derived from the intron motif. This option changes the output alignments: reads with inconsistent and/or non-canonical introns are filtered out.

`--outSAMattributes`

default: `Standard`

string(s): a string of desired SAM attributes, in the order desired for the output SAM. Tags can be listed in any combination/order.

***Presets:

`None`
: no attributes

`Standard`
: NH HI AS nM

`All`
: NH HI AS nM NM MD jM jI MC ch

***Alignment:

NH
: number of loci the reads maps to: =1 for unique mappers, >1 for multimappers. Standard SAM tag.

HI
: multiple alignment index, starts with –outSAMattrIHstart (=1 by default). Standard SAM tag.

AS
: local alignment score, +1/-1 for matches/mismateches, score* penalties for indels and gaps. For PE reads, total score for two mates. Stadnard SAM tag.

nM
: number of mismatches. For PE reads, sum over two mates.

NM
: edit distance to the reference (number of mismatched + inserted + deleted bases) for each mate. Standard SAM tag.

MD
: string encoding mismatched and deleted reference bases (see standard SAM specifications). Standard SAM tag.

jM
: intron motifs for all junctions (i.e. N in CIGAR): 0: non-canonical; 1: GT/AG, 2: CT/AC, 3: GC/AG, 4: CT/GC, 5: AT/AC, 6: GT/AT. If splice junctions database is used, and a junction is annotated, 20 is added to its motif value.

jI
: start and end of introns for all junctions (1-based).

XS
: alignment strand according to –outSAMstrandField.

MC
: mate's CIGAR string. Standard SAM tag.

ch
: marks all segment of all chimeric alingments for –chimOutType WithinBAM output.

cN
: number of bases clipped from the read ends: 5' and 3'

***Variation:

vA
: variant allele

vG
: genomic coordinate of the variant overlapped by the read.

vW
: 1 - alignment passes WASP filtering; 2,3,4,5,6,7 - alignment does not pass WASP filtering. Requires –waspOutputMode SAMtag.

ha
: haplotype (1/2) when mapping to the diploid genome. Requires genome generated with –genomeTransformType Diploid .

***STARsolo:

`CR CY UR UY`
: sequences and quality scores of cell barcodes and UMIs for the solo* demultiplexing.

`GX GN`
: gene ID and gene name for unique-gene reads.

`gx gn`
: gene IDs and gene names for unique- and multi-gene reads.

`CB UB`
: error-corrected cell barcodes and UMIs for solo* demultiplexing. Requires –outSAMtype BAM SortedByCoordinate.

`sM`
: assessment of CB and UMI.

`sS`
: sequence of the entire barcode (CB,UMI,adapter).

`sQ`
: quality of the entire barcode.

`sF`
: type of feature overlap and number of features for each alignment

***Unsupported/undocumented:

`rB`
: alignment block read/genomic coordinates.

`vR`
: read coordinate of the variant.

`--outSAMattrIHstart`

default: `1`

int>=0: start value for the IH attribute. 0 may be required by some downstream software, such as Cufflinks or StringTie.

`--outSAMunmapped`

default: `None`

string(s): output of unmapped reads in the SAM format

1st word:

`None`
: no output

`Within`
: output unmapped reads within the main SAM file (i.e. Aligned.out.sam)

2nd word:

`KeepPairs`
: record unmapped mate for each alignment, and, in case of unsorted output, keep it adjacent to its mapped mate. Only affects multi-mapping reads.

`--outSAMorder`

default: `Paired`

string: type of sorting for the SAM output

Paired: one mate after the other for all paired alignments

PairedKeepInputOrder: one mate after the other for all paired alignments, the order is kept the same as in the input FASTQ files

`--outSAMprimaryFlag`

default: `OneBestScore`

string: which alignments are considered primary - all others will be marked with 0x100 bit in the FLAG

`OneBestScore`
: only one alignment with the best score is primary

`AllBestScore`
: all alignments with the best score are primary

`--outSAMreadID`

default: `Standard`

string: read ID record type

`Standard`
: first word (until space) from the FASTx read ID line, removing /1,/2 from the end

`Number`
: read number (index) in the FASTx file

`--outSAMmapqUnique`

default: `255`

int: 0 to 255: the MAPQ value for unique mappers

`--outSAMflagOR`

default: `0`

int: 0 to 65535: sam FLAG will be bitwise OR'd with this value, i.e. FLAG=FLAG — outSAMflagOR. This is applied after all flags have been set by STAR, and after outSAMflagAND. Can be used to set specific bits that are not set otherwise.

`--outSAMflagAND`

default: `65535`

int: 0 to 65535: sam FLAG will be bitwise AND'd with this value, i.e. FLAG=FLAG & outSAMflagOR. This is applied after all flags have been set by STAR, but before outSAMflagOR. Can be used to unset specific bits that are not set otherwise.

`--outSAMattrRGline`

default: `-`

string(s): SAM/BAM read group line. The first word contains the read group identifier and must start with "ID:", e.g. –outSAMattrRGline ID:xxx CN:yy "DS:z z z".

xxx will be added as RG tag to each output alignment. Any spaces in the tag values have to be double quoted.

Comma separated RG lines correspons to different (comma separated) input files in –readFilesIn. Commas have to be surrounded by spaces, e.g.

–outSAMattrRGline ID:xxx , ID:zzz "DS:z z" , ID:yyy DS:yyyy

`--outSAMheaderHD`

default: `-`

strings: @HD (header) line of the SAM header

`--outSAMheaderPG`

default: `-`

strings: extra @PG (software) line of the SAM header (in addition to STAR)

`--outSAMheaderCommentFile`

default: `-`

string: path to the file with @CO (comment) lines of the SAM header

`--outSAMfilter`

default: `None`

string(s): filter the output into main SAM/BAM files

`KeepOnlyAddedReferences`
: only keep the reads for which all alignments are to the extra reference sequences added with –genomeFastaFiles at the mapping stage.

`KeepAllAddedReferences`
: keep all alignments to the extra reference sequences added with –genomeFastaFiles at the mapping stage.

`--outSAMmultNmax`

default: `-1`

int: max number of multiple alignments for a read that will be output to the SAM/BAM files. Note that if this value is not equal to -1, the top scoring alignment will be output first

`-1`
: all alignments (up to –outFilterMultimapNmax) will be output

`--outSAMtlen`

default: `1`

int: calculation method for the TLEN field in the SAM/BAM files

`1`
: leftmost base of the (+)strand mate to rightmost base of the (-)mate. (+)sign for the (+)strand mate

`2`
: leftmost base of any mate to rightmost base of any mate. (+)sign for the mate with the leftmost base. This is different from 1 for overlapping mates with protruding ends

`--outBAMcompression`

default: `1`

int: -1 to 10 BAM compression level, -1=default compression (6?), 0=no compression, 10=maximum compression

`--outBAMsortingThreadN`

default: `0`

int: >=0: number of threads for BAM sorting. 0 will default to min(6,–runThreadN).

`--outBAMsortingBinsN`

default: `50`

int: >0: number of genome bins for coordinate-sorting

#### 17.14 BAM processing

`--bamRemoveDuplicatesType`

default: `-`

string: mark duplicates in the BAM file, for now only works with (i) sorted BAM fed with inputBAMfile, and (ii) for paired-end alignments only

`-`
: no duplicate removal/marking

`UniqueIdentical`
: mark all multimappers, and duplicate unique mappers. The coordinates, FLAG, CIGAR must be identical

`UniqueIdenticalNotMulti`
: mark duplicate unique mappers but not multimappers.

`--bamRemoveDuplicatesMate2basesN`

default: `0`

int>0: number of bases from the 5' of mate 2 to use in collapsing (e.g. for RAMPAGE)

#### 17.15 Output Wiggle

`--outWigType`

default: `None`

string(s): type of signal output, e.g. "bedGraph" OR "bedGraph read1_5p". Requires sorted BAM: –outSAMtype BAM SortedByCoordinate .

1st word:

`None`
: no signal output

`bedGraph`
: bedGraph format

`wiggle`
: wiggle format

2nd word:

`read1_5p`
: signal from only 5' of the 1st read, useful for CAGE/RAMPAGE etc

`read2`
: signal from only 2nd read

`--outWigStrand`

default: `Stranded`

string: strandedness of wiggle/bedGraph output

`Stranded`
: separate strands, str1 and str2

`Unstranded`
: collapsed strands

`--outWigReferencesPrefix`

default: `-`

string: prefix matching reference names to include in the output wiggle file, e.g. ”chr”, default ”-” - include all references

`--outWigNorm`

default: `RPM`

string: type of normalization for the signal

`RPM`
: reads per million of mapped reads

`None`
: no normalization, ”raw” counts

#### 17.16 Output Filtering

`--outFilterType`

default: `Normal`

string: type of filtering

`Normal`
: standard filtering using only current alignment

`BySJout`
: keep only those reads that contain junctions that passed filtering into SJ.out.tab

`--outFilterMultimapScoreRange`

default: `1`

int: the score range below the maximum score for multimapping alignments

`--outFilterMultimapNmax`

default: `10`

int: maximum number of loci the read is allowed to map to. Alignments (all of them) will be output only if the read maps to no more loci than this value.

Otherwise no alignments will be output, and the read will be counted as "mapped to too many loci" in the Log.final.out .

`--outFilterMismatchNmax`

default: `10`

int: alignment will be output only if it has no more mismatches than this value.

`--outFilterMismatchNoverLmax`

default: `0.3`

real: alignment will be output only if its ratio of mismatches to *mapped* length is less than or equal to this value.

`--outFilterMismatchNoverReadLmax`

default: `1.0`

real: alignment will be output only if its ratio of mismatches to *read* length is less than or equal to this value.

`--outFilterScoreMin`

default: `0`

int: alignment will be output only if its score is higher than or equal to this value.

`--outFilterScoreMinOverLread`

default: `0.66`

real: same as outFilterScoreMin, but normalized to read length (sum of mates' lengths for paired-end reads)

`--outFilterMatchNmin`

default: `0`

int: alignment will be output only if the number of matched bases is higher than or equal to this value.

`--outFilterMatchNminOverLread`

default: `0.66`

real: sam as outFilterMatchNmin, but normalized to the read length (sum of mates' lengths for paired-end reads).

`--outFilterIntronMotifs`

default: `None`

string: filter alignment using their motifs

`None`
: no filtering

`RemoveNoncanonical`
: filter out alignments that contain non-canonical junctions

`RemoveNoncanonicalUnannotated`
: filter out alignments that contain non-canonical unannotated junctions when using annotated splice junctions database. The annotated non-canonical junctions will be kept.

`--outFilterIntronStrands`

default: `RemoveInconsistentStrands`

string: filter alignments

`RemoveInconsistentStrands`
: remove alignments that have junctions with inconsistent strands

`None`
: no filtering

#### 17.17 Output splice junctions (SJ.out.tab)

`--outSJtype`

default: `Standard`

string: type of splice junction output

`Standard`
: standard SJ.out.tab output

`None`
: no splice junction output

#### 17.18 Output Filtering: Splice Junctions

`--outSJfilterReads`

default: `All`

string: which reads to consider for collapsed splice junctions output

`All`
: all reads, unique- and multi-mappers

`Unique`
: uniquely mapping reads only

`--outSJfilterOverhangMin`

default: `30 12 12 12`

4 integers: minimum overhang length for splice junctions on both sides for: (1) non-canonical motifs, (2) GT/AG and CT/AC motif, (3) GC/AG and CT/GC motif, (4) AT/AC and GT/AT motif. -1 means no output for that motif

does not apply to annotated junctions

`--outSJfilterCountUniqueMin`

default: `3 1 1 1`

4 integers: minimum uniquely mapping read count per junction for: (1) non-canonical motifs, (2) GT/AG and CT/AC motif, (3) GC/AG and CT/GC motif, (4) AT/AC and GT/AT motif. -1 means no output for that motif

Junctions are output if one of outSJfilterCountUniqueMin OR outSJfilterCountTotalMin conditions are satisfied

does not apply to annotated junctions

`--outSJfilterCountTotalMin`

default: `3 1 1 1`

4 integers: minimum total (multi-mapping+unique) read count per junction for: (1) non-canonical motifs, (2) GT/AG and CT/AC motif, (3) GC/AG and CT/GC motif, (4) AT/AC and GT/AT motif. -1 means no output for that motif

Junctions are output if one of outSJfilterCountUniqueMin OR outSJfilterCountTotalMin conditions are satisfied

does not apply to annotated junctions

`--outSJfilterDistToOtherSJmin`

default: `10 0 5 10`

4 integers>=0: minimum allowed distance to other junctions' donor/acceptor

does not apply to annotated junctions

`--outSJfilterIntronMaxVsReadN`

default: `50000 100000 200000`

N integers>=0: maximum gap allowed for junctions supported by 1,2,3,,,N reads

i.e. by default junctions supported by 1 read can have gaps <=50000b, by 2 reads: <=100000b, by 3 reads: <=200000. by >=4 reads any gap <=alignIntronMax

does not apply to annotated junctions

##### 17.19 Scoring

--scoreGap

default: 0

int: splice junction penalty (independent on intron motif)

--scoreGapNoncan

default: -8

int: non-canonical junction penalty (in addition to scoreGap)

--scoreGapGCAG

default: -4

int: GC/AG and CT/GC junction penalty (in addition to scoreGap)

--scoreGapATAC

default: -8

int: AT/AC and GT/AT junction penalty (in addition to scoreGap)

--scoreGenomicLengthLog2scale

default: -0.25

int: extra score logarithmically scaled with genomic length of the alignment: scoreGenomicLengthLog2scale*log2(genomicLength)

--scoreDelOpen

default: -2

int: deletion open penalty

--scoreDelBase

default: -2

int: deletion extension penalty per base (in addition to scoreDelOpen)

--scoreInsOpen

default: -2

int: insertion open penalty

--scoreInsBase

default: -2

int: insertion extension penalty per base (in addition to scoreInsOpen)

--scoreStitchSJshift

default: 1

int: maximum score reduction while searching for SJ boundaries in the stitching step

#### 17.20 Alignments and Seeding

`--seedSearchStartLmax`

default: 50

int>0: defines the search start point through the read - the read is split into pieces no longer than this value

`--seedSearchStartLmaxOverLread`

default: 1.0

real: seedSearchStartLmax normalized to read length (sum of mates' lengths for paired-end reads)

`--seedSearchLmax`

default: 0

int>=0: defines the maximum length of the seeds, if =0 seed length is not limited

`--seedMultimapNmax`

default: 10000

int>0: only pieces that map fewer than this value are utilized in the stitching procedure

`--seedPerReadNmax`

default: 1000

int>0: max number of seeds per read

`--seedPerWindowNmax`

default: 50

int>0: max number of seeds per window

`--seedNoneLociPerWindow`

default: 10

int>0: max number of one seed loci per window

`--seedSplitMin`

default: 12

int>0: min length of the seed sequences split by Ns or mate gap

--seedMapMin

default: 5

int>0: min length of seeds to be mapped

--alignIntronMin

default: 21

int: minimum intron size, genomic gap is considered intron if its length>=alignIntronMin, otherwise it is considered Deletion

--alignIntronMax

default: 0

int: maximum intron size, if 0, max intron size will be determined by (2^winBinNbits)*winAnchorDistNbins

--alignMatesGapMax

default: 0

int: maximum gap between two mates, if 0, max intron gap will be determined by (2^winBinNbits)*winAnchorDistNbins

--alignSJoverhangMin

default: 5

int>0: minimum overhang (i.e. block size) for spliced alignments

--alignSJstitchMismatchNmax

default: 0 -1 0 0

4*int>=0: maximum number of mismatches for stitching of the splice junctions (-1: no limit).

(1) non-canonical motifs, (2) GT/AG and CT/AC motif, (3) GC/AG and CT/GC motif, (4) AT/AC and GT/AT motif.

--alignSJDBoverhangMin

default: 3

int>0: minimum overhang (i.e. block size) for annotated (sjdb) spliced alignments

--alignSplicedMateMapLmin

default: 0

int>0: minimum mapped length for a read mate that is spliced

`--alignSplicedMateMapLminOverLmate`

default: `0.66`

real>0: alignSplicedMateMapLmin normalized to mate length

`--alignWindowsPerReadNmax`

default: `10000`

int>0: max number of windows per read

`--alignTranscriptsPerWindowNmax`

default: `100`

int>0: max number of transcripts per window

`--alignTranscriptsPerReadNmax`

default: `10000`

int>0: max number of different alignments per read to consider

`--alignEndsType`

default: `Local`

string: type of read ends alignment

`Local`
: standard local alignment with soft-clipping allowed

`EndToEnd`
: force end-to-end read alignment, do not soft-clip

`Extend5pOfRead1`
: fully extend only the 5p of the read1, all other ends: local alignment

`Extend5pOfReads12`
: fully extend only the 5p of the both read1 and read2, all other ends: local alignment

`--alignEndsProtrude`

default: `0 ConcordantPair`

int, string: allow protrusion of alignment ends, i.e. start (end) of the +strand mate downstream of the start (end) of the -strand mate

1st word: int: maximum number of protrusion bases allowed

2nd word: string:

`ConcordantPair`
: report alignments with non-zero protrusion as concordant pairs

`DiscordantPair`
: report alignments with non-zero protrusion as discordant pairs

`--alignSoftClipAtReferenceEnds`

default: `Yes`

string: allow the soft-clipping of the alignments past the end of the chromosomes

`Yes`
: allow

`No`
: prohibit, useful for compatibility with Cufflinks

`--alignInsertionFlush`

default: `None`

string: how to flush ambiguous insertion positions

`None`
: insertions are not flushed

`Right`
: insertions are flushed to the right

##### 17.21 Paired-End reads

`--peOverlapNbasesMin`

default: `0`

int>=0: minimum number of overlapping bases to trigger mates merging and realignment. Specify >0 value to switch on the "merginf of overlapping mates" algorithm.

`--peOverlapMMp`

default: `0.01`

real, >=0 & <1: maximum proportion of mismatched bases in the overlap area

#### 17.22 Windows, Anchors, Binning

`--winAnchorMultimapNmax`

default: `50`

int>0: max number of loci anchors are allowed to map to

`--winBinNbits`

default: `16`

int>0: =log2(winBin), where winBin is the size of the bin for the windows/clustering, each window will occupy an integer number of bins.

`--winAnchorDistNbins`

default: `9`

int>0: max number of bins between two anchors that allows aggregation of anchors into one window

`--winFlankNbins`

default: `4`

int>0: log2(winFlank), where win Flank is the size of the left and right flanking regions for each window

`--winReadCoverageRelativeMin`

default: `0.5`

real>=0: minimum relative coverage of the read sequence by the seeds in a window, for STARlong algorithm only.

`--winReadCoverageBasesMin`

default: `0`

int>0: minimum number of bases covered by the seeds in a window , for STARlong algorithm only.

#### 17.23 Chimeric Alignments

--chimOutType

default: Junctions

string(s): type of chimeric output

Junctions
: Chimeric.out.junction

SeparateSAMold
: output old SAM into separate Chimeric.out.sam file

WithinBAM
: output into main aligned BAM files (Aligned.*.bam)

WithinBAM HardClip
: (default) hard-clipping in the CIGAR for supplemental chimeric alignments (default if no 2nd word is present)

WithinBAM SoftClip
: soft-clipping in the CIGAR for supplemental chimeric alignments

--chimSegmentMin

default: 0

int>=0: minimum length of chimeric segment length, if ==0, no chimeric output

--chimScoreMin

default: 0

int>=0: minimum total (summed) score of the chimeric segments

--chimScoreDropMax

default: 20

int>=0: max drop (difference) of chimeric score (the sum of scores of all chimeric segments) from the read length

--chimScoreSeparation

default: 10

int>=0: minimum difference (separation) between the best chimeric score and the next one

--chimScoreJunctionNonGTAG

default: -1

int: penalty for a non-GT/AG chimeric junction

`--chimJunctionOverhangMin`

default: `20`

int>=0: minimum overhang for a chimeric junction

`--chimSegmentReadGapMax`

default: `0`

int>=0: maximum gap in the read sequence between chimeric segments

`--chimFilter`

default: `banGenomicN`

string(s): different filters for chimeric alignments

`None`
: no filtering

`banGenomicN`
: Ns are not allowed in the genome sequence around the chimeric junction

`--chimMainSegmentMultNmax`

default: `10`

int>=1: maximum number of multi-alignments for the main chimeric segment. =1 will prohibit multimapping main segments.

`--chimMultimapNmax`

default: `0`

int>=0: maximum number of chimeric multi-alignments

`0`
: use the old scheme for chimeric detection which only considered unique alignments

`--chimMultimapScoreRange`

default: `1`

int>=0: the score range for multi-mapping chimeras below the best chimeric score. Only works with –chimMultimapNmax > 1

`--chimNonchimScoreDropMin`

default: `20`

int>=0: to trigger chimeric detection, the drop in the best non-chimeric alignment score with respect to the read length has to be greater than this value

--chimOutJunctionFormat

default: 0

int: formatting type for the Chimeric.out.junction file

0
: no comment lines/headers

1
: comment lines at the end of the file: command line and Nreads: total, unique/multi-mapping

#### 17.24 Quantification of Annotations

--quantMode

default: -

string(s): types of quantification requested

-
: none

TranscriptomeSAM
: output SAM/BAM alignments to transcriptome into a separate file

GeneCounts
: count reads per gene

--quantTranscriptomeBAMcompression

default: 1

int: -2 to 10 transcriptome BAM compression level

-2
: no BAM output

-1
: default compression (6?)

0
: no compression

10
: maximum compression

--quantTranscriptomeSAMoutput

default: BanSingleEnd_BanIndels_ExtendSoftclip

string: alignment filtering for TranscriptomeSAM output

BanSingleEnd_BanIndels_ExtendSoftclip
prohibit indels and single-end alignments, extend softclips - compatible with RSEM

BanSingleEnd
prohibit single-end alignments, allow indels and softclips

BanSingleEnd_ExtendSoftclip
prohibit single-end alignments, extend softclips, allow indels

#### 17.25 2-pass Mapping

--twopassMode

default: None

string: 2-pass mapping mode.

None
1-pass mapping

Basic
basic 2-pass mapping, with all 1st pass junctions inserted into the genome indices on the fly

--twopass1readsN

default: -1

int: number of reads to process for the 1st step. Use very large number (or default -1) to map all reads in the first step.

#### 17.26 WASP parameters

--waspOutputMode

default: None

string: WASP allele-specific output type. This is re-implementation of the original WASP mappability filtering by Bryce van de Geijn, Graham McVicker, Yoav Gilad & Jonathan K Pritchard. Please cite the original WASP paper: Nature Methods 12, 1061–1063 (2015), https://www.nature.com/articles/nmeth.3582 .

SAMtag
add WASP tags to the alignments that pass WASP filtering

#### 17.27 STARsolo (single cell RNA-seq) parameters

--soloType

default: None

string(s): type of single-cell RNA-seq

CB_UMI_Simple
(a.k.a. Droplet) one UMI and one Cell Barcode of fixed length in read2, e.g. Drop-seq and 10X Chromium.

CB_UMI_Complex
multiple Cell Barcodes of varying length, one UMI of fixed length and one adapter sequence of fixed length are allowed in read2 only (e.g. inDrop, ddSeq).

CB_samTagOut
output Cell Barcode as CR and/or CB SAm tag. No UMI counting. –readFilesIn cDNA_read1 [cDNA_read2 if paired-end] CellBarcode_read . Requires –outSAMtype BAM Unsorted [and/or SortedByCoordinate]

SmartSeq
Smart-seq: each cell in a separate FASTQ (paired- or single-end), barcodes are corresponding read-groups, no UMI sequences, alignments deduplicated according to alignment start and end (after extending soft-clipped bases)

--soloCBtype

default: Sequence

string: cell barcode type

Sequence: cell barcode is a sequence (standard option)

String: cell barcode is an arbitrary string

--soloCBwhitelist

default: -

string(s): file(s) with whitelist(s) of cell barcodes. Only –soloType CB_UMI_Complex allows more than one whitelist file.

None
no whitelist: all cell barcodes are allowed

--soloCBstart

default: 1

int>0: cell barcode start base

`--soloCBlen`

default: `16`

int>0: cell barcode length

`--soloUMIstart`

default: `17`

int>0: UMI start base

`--soloUMIlen`

default: `10`

int>0: UMI length

`--soloBarcodeReadLength`

default: `1`

int: length of the barcode read

`1`
: equal to sum of soloCBlen+soloUMIlen

`0`
: not defined, do not check

`--soloBarcodeMate`

default: `0`

int: identifies which read mate contains the barcode (CB+UMI) sequence

`0`
: barcode sequence is on separate read, which should always be the last file in the –readFilesIn listed

`1`
: barcode sequence is a part of mate 1

`2`
: barcode sequence is a part of mate 2

`--soloCBposition`

default: `-`

strings(s): position of Cell Barcode(s) on the barcode read.

Presently only works with –soloType CB_UMI_Complex, and barcodes are assumed to be on Read2.

Format for each barcode: startAnchor_startPosition_endAnchor_endPosition

start(end)Anchor defines the Anchor Base for the CB: 0: read start; 1: read end; 2: adapter start; 3: adapter end

start(end)Position is the 0-based position with of the CB start(end) with respect to the Anchor Base

String for different barcodes are separated by space.

Example: inDrop (Zilionis et al, Nat. Protocols, 2017):

–soloCBposition 0_0_2_-1 3_1_3_8

`--soloUMIposition`

default: `-`

string: position of the UMI on the barcode read, same as soloCBposition

Example: inDrop (Zilionis et al, Nat. Protocols, 2017):

–soloCBposition 3_9_3_14

`--soloAdapterSequence`

default: `-`

string: adapter sequence to anchor barcodes. Only one adapter sequence is allowed.

`--soloAdapterMismatchesNmax`

default: `1`

int>0: maximum number of mismatches allowed in adapter sequence.

`--soloCBmatchWLtype`

default: `1MM_multi`

string: matching the Cell Barcodes to the WhiteList

`Exact`
: only exact matches allowed

`1MM`
: only one match in whitelist with 1 mismatched base allowed. Allowed CBs have to have at least one read with exact match.

`1MM_multi`
: multiple matches in whitelist with 1 mismatched base allowed, posterior probability calculation is used choose one of the matches.

Allowed CBs have to have at least one read with exact match. This option matches best with CellRanger 2.2.0

1MM_multi_pseudocounts
: same as 1MM_Multi, but pseudocounts of 1 are added to all whitelist barcodes.

1MM_multi_Nbase_pseudocounts
: same as 1MM_multi_pseudocounts, multimatching to WL is allowed for CBs with N-bases. This option matches best with CellRanger >= 3.0.0

EditDist_2
: allow up to edit distance of 3 fpr each of the barcodes. May include one deletion + one insertion. Only works with –soloType CB_UMI_Complex. Matches to multiple passlist barcdoes are not allowed. Similar to ParseBio Split-seq pipeline.

--soloInputSAMattrBarcodeSeq

default: -

string(s): when inputting reads from a SAM file (–readsFileType SAM SE/PE), these SAM attributes mark the barcode sequence (in proper order).

For instance, for 10X CellRanger or STARsolo BAMs, use –soloInputSAMattrBarcodeSeq CR UR .

This parameter is required when running STARsolo with input from SAM.

--soloInputSAMattrBarcodeQual

default: -

string(s): when inputting reads from a SAM file (–readsFileType SAM SE/PE), these SAM attributes mark the barcode qualities (in proper order).

For instance, for 10X CellRanger or STARsolo BAMs, use –soloInputSAMattrBarcodeQual CY UY .

If this parameter is '-' (default), the quality 'H' will be assigned to all bases.

--soloStrand

default: Forward

string: strandedness of the solo libraries:

Unstranded
: no strand information

Forward
: read strand same as the original RNA molecule

Reverse
: read strand opposite to the original RNA molecule

`--soloFeatures`

default: `Gene`

string(s): genomic features for which the UMI counts per Cell Barcode are collected

`Gene`
: genes: reads match the gene transcript

`SJ`
: splice junctions: reported in SJ.out.tab

`GeneFull`
: full gene (pre-mRNA): count all reads overlapping genes' exons and introns

`GeneFull_ExonOverIntron`
: full gene (pre-mRNA): count all reads overlapping genes' exons and introns: prioritize 100% overlap with exons

`GeneFull_Ex50pAS`
: full gene (pre-RNA): count all reads overlapping genes' exons and introns: prioritize >50% overlap with exons. Do not count reads with 100% exonic overlap in the antisense direction.

`--soloMultiMappers`

default: `Unique`

string(s): counting method for reads mapping to multiple genes

`Unique`
: count only reads that map to unique genes

`Uniform`
: uniformly distribute multi-genic UMIs to all genes

`Rescue`
: distribute UMIs proportionally to unique+uniform counts ( first iteration of EM)

`PropUnique`
: distribute UMIs proportionally to unique mappers, if present, and uniformly if not.

`EM`
: multi-gene UMIs are distributed using Expectation Maximization algorithm

`--soloUMIdedup`

default: `1MM_All`

string(s): type of UMI deduplication (collapsing) algorithm

`1MM_All`
: all UMIs with 1 mismatch distance to each other are collapsed (i.e. counted once).

`1MM_Directional_UMItools`
: follows the "directional" method from the UMI-tools by Smith, Heger and Sudbery (Genome Research 2017).

`1MM_Directional`
: same as 1MM_Directional_UMItools, but with more stringent criteria for duplicate UMIs

`Exact`
: only exactly matching UMIs are collapsed.

`NoDedup`
: no deduplication of UMIs, count all reads.

`1MM_CR`
: CellRanger2-4 algorithm for 1MM UMI collapsing.

`--soloUMIfiltering`

default: `-`

string(s): type of UMI filtering (for reads uniquely mapping to genes)

`-`
: basic filtering: remove UMIs with N and homopolymers (similar to CellRanger 2.2.0).

`MultiGeneUMI`
: basic + remove lower-count UMIs that map to more than one gene.

`MultiGeneUMI_All`
: basic + remove all UMIs that map to more than one gene.

`MultiGeneUMI_CR`
: basic + remove lower-count UMIs that map to more than one gene, matching CellRanger > 3.0.0 .

Only works with –soloUMIdedup 1MM_CR

`--soloOutFileNames`

default: `Solo.out/ features.tsv barcodes.tsv matrix.mtx`

string(s): file names for STARsolo output:

file_name_prefix gene_names barcode_sequences cell_feature_count_matrix

`--soloCellFilter`

default: `CellRanger2.2 3000 0.99 10`

string(s): cell filtering type and parameters

None
: do not output filtered cells

TopCells
: only report top cells by UMI count, followed by the exact number of cells

CellRanger2.2
: simple filtering of CellRanger 2.2.

Can be followed by numbers: number of expected cells, robust maximum percentile for UMI count, maximum to minimum ratio for UMI count

The harcoded values are from CellRanger: nExpectedCells=3000; maxPercentile=0.99; maxMinRatio=10

EmptyDrops_CR
: EmptyDrops filtering in CellRanger flavor. Please cite the original EmptyDrops paper: A.T.L Lun et al, Genome Biology, 20, 63 (2019): https://genomebiology.biomedcentral.com/articles/10.1186/s13059-019-1662-y

Can be followed by 10 numeric parameters: nExpectedCells maxPercentile maxMinRatio indMin indMax umiMin umiMinFracMedian candMaxN FDR simN

The harcoded values are from CellRanger: 3000 0.99 10 45000 90000 500 0.01 20000 0.01 10000

`--soloOutFormatFeaturesGeneField3`

default: "Gene Expression"

string(s): field 3 in the Gene features.tsv file. If "-", then no 3rd field is output.

`--soloCellReadStats`

default: None

string: Output reads statistics for each CB

Standard
: standard output
